# Postnatal sensorimotor experience causally shapes hemispheric specialization across motor and language networks

**DOI:** 10.64898/2026.09.22.753435

**Authors:** Yuqi Liu, Jiayu Sun, Lianzheng Zhao, Liang Chen, Liping Wang

## Abstract

The functional organization of the human brain is shaped by the interplay between innate constraints and postnatal experience. Hemispheric specialization, as a fundamental organizing principle of brain function, raises unresolved questions about how postnatal experience shapes lateralization and importantly, whether different lateralized cognitive systems develop through shared or independent mechanisms. We leveraged a natural experiment in individuals with obstetric brachial plexus injury (OBPI) in the right upper limb, resulting in lifelong left-hand dominance dissociated from the typical predisposition, to address these questions. Using functional MRI experiments during manual motor tasks, we found that altered hand-use experience selectively reshaped hemispheric organization in higher-order frontoparietal motor regions. Specifically, individuals with right OBPI showed reduced left-hemisphere dominance and rightward reorganization of abstract, hand-independent action representations relative to controls and individuals with left-side OBPI, whereas primary sensorimotor and subcortical regions preserved their canonical lateralizedl organization. Strikingly, experience-dependent motor reorganization extended beyond the motor system individuals with right OBPI exhibited reduced left lateralization of the language network in Broca’s area (BA44) during sentence listening. Crucially, language lateralization in BA44 covaried with the left lateralization of frontal motor areas, particularly for the compensatory, functional left hand. These findings demonstrate that postnatal bodily experience shapes the developmental emergence of hemispheric specialization, specifically in higher-order motor and language networks, and revealed coordinated organization between action and language systems.

## Introduction

The functional organization of human cortex emerges through interactions between intrinsic constraints and postnatal experience. One of the fundamental organization principles is hemispheric specialization, which follows remarkably systematic patterns across cognitive domains. For example, language and spatial attention are reliably dominated by the opposite hemispheres (Cai et al., 2013; Toga & Thompson, 2003; Knecht et al., 2000; Corbetta & Schulman, 2011). Such regularities suggest that the hemispheric organization of different cognitive systems may not emerge independently, but instead reflect common constraints that coordinate their distribution (Gotts et al., 2013; Toga & Thompson, 2003). However, the mechanisms that give rise to these coordinated patterns remain poorly understood. Two fundamental questions remain unresolved: to what extent is hemispheric specialization shaped by postnatal experience rather than innate predisposition, and do different lateralized cognitive systems develop independently or through shared mechanisms?

These questions have been challenging to address because the mapping between experience and functional lateralization is usually indirect. Among lateralized cognitive systems, the motor system provides a particularly tractable model because asymmetries can be readily characterized at both behavioral and neural levels. Whereas the primary sensorimotor cortices show contralateral dominance (Penfield & Boldrey, 1937; Yousry et al., 1997), higher-order frontoparietal areas display left lateralization for reaching and grasping (Di Caro et al., 2025; Merrick et al., 2022; Kumar et al., 2020) and tool use (Thibault et al., 2021; Buxbaum et al., 2014), regardless of which hand is executing the action. These findings suggest that higher-order motor areas in the left hemisphere dominate in encoding abstract action information independent of hand. Notably, the degree of left lateralization is reduced in left-handers (Vingerhoets et al., 2012; Kroliczak et al., 2021), demonstrating statistical correlation between hemispheric specialization and hand use experience. However, because innate predispositions and postnatal experience are inseparable under natural conditions, existing evidence cannot disentangle whether asymmetric hand use causally contributes to hemispheric dominance, or whether both handedness and hemispheric dominance are innately determined.

Experience-dependent reorganization also provides a unique opportunity to test whether different lateralized cognitive systems share common developmental constraints. Beyond the motor system, language is typically lateralized to the left hemisphere in right-handers and more variable in left-handers (Knecht et al., 2000; Wiberg et al., 2019; Malik-Moraleda et al., 2022), showing similar contingency to handedness as with the higher-order motor system. Multiple lines of evidence suggest close links between action and language systems. At the behavioral level, the development of manual dexterity is associated with language abilities in children (Iverson, 2010; Contino et al., 2025; Tseng & Hsu, 2025), while training on precision tool use and syntactic processing showed mutual facilitation in adults, suggesting shared underlying mechanisms (Thibault et al., 2021). At the neural level, both systems follow a transition from relatively bilateral organization in childhood to increasingly left-lateralized patterns in adulthood (Olulade et al., 2020; Ozernov-Palchik et al., 2026; Biagi et al., 2015; Dehaene-Lambertz et al., 2006), and the degree of left lateralization is correlated between motor and language networks in resting state functional connectivity (Gotts et al., 2013). Together, these findings raise the possibility that motor and language emerge through shared developmental processes rather than independent maturation. One influential proposal is that action and language rely on common abstract computations, particularly hierarchical sequencing operations implemented by overlapping frontal and subcortical circuits (Greenfield, 1991; Koechlin & Jubault, 2006; Thibault et al., 2021; Wandelt et al., 2022; Friederici, 2023). If hemispheric specialization is governed by shared developmental mechanisms across cognitive domains, experience-dependent reorganization of the motor system should propagate beyond the motor domain and reshape the lateralization of language.

Addressing these questions necessitates the dissociation between innate predisposition of hemispheric specialization and postnatal experience. We leveraged a unique population—individuals with obstetric brachial plexus injuries (OBPI)—to investigate both the developmental origins of hemispheric specialization and the relationship between action and language systems . In severe cases of OBPI, peripheral nerves connecting the spinal cord and upper limb are severely injured during birth, resulting in lifelong motor impairment of the affected arm and altered sensorimotor experience throughout postnatal development (Malessy et al., 2009). Critically, individuals with right-side OBPI (R-OBPI) are compelled to rely predominantly on their left hand from birth onward, despite an innate predisposition for left-hemisphere dominance. This dissociation makes OBPI an exceptional natural experiment for testing causal effects of sensorimotor experience on hemispheric specialization (**Fig. 1A**).

**Figure 1.**
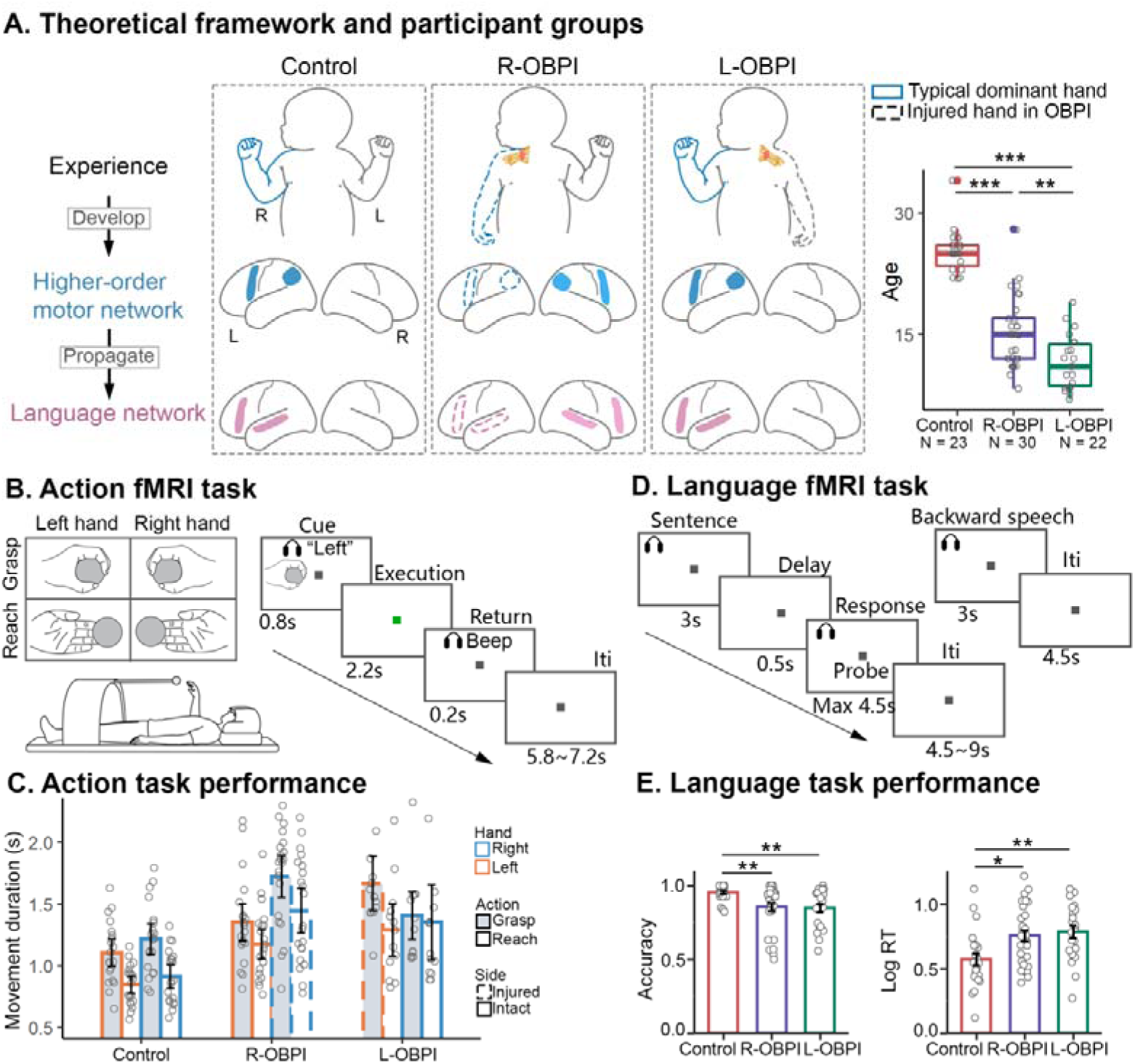
Theoretical framework and experimental design of the current study. **(A)** Control participants, individuals with right obstetric brachial plexus injury (R-OBPI), and individuals with left obstetric brachial plexus injury (L-OBPI) were tested. Blue outlines indicate the functionally dominant hand, whereas dashed outlines denote the injured upper limb in the OBPI groups. The conceptual framework proposes that asymmetric hand use during development shapes higher-order motor networks, which in turn influences the lateralization of the language network. Box plots show the age distribution of each group. (B)A 2 (Grasping, Reaching) by 2 (Left hand, Right hand) design was adopted in the action fMRI task. At trial onset, an auditory cue (“left” or “right”) and a visual instruction depicting the action type was presented for 800 ms. Participants then executed the instructed action toward a centrally located object until a 200-ms auditory beep signaling return to the home position, followed by inter-trial interval (iti). (C)Movement duration during the action fMRI task. As expected, the affected side in individuals with OBPI performed slower than the unaffected side (post-hoc *p*s < .003). Across hands and groups, grasping action required a longer time to perform than reaching (all post-hoc *p*s < .001, except for the right hand in L-OBPI for which *p* = .061). **(D)** Participants listened to spoken sentences (e.g. “A monkey is riding on an elephant and picking apples.”) or backward speech during the language task. Following each sentence, a probe sentence (e.g. “The monkey is picking apples.”) was presented both auditorily and visually, and participants judged whether the two sentences were semantically consistent by pressing one of two buttons. (E) Accuracy and reaction time of the sentence judgment task. The control group performed more accurately (*p*s < .002, with *p* = .919 between L-OBPI and R-OBPI), as well as faster than the OBPI groups (*p*s < .015, with *p* = .882 between L-OBPI and R-OBPI).

If hemispheric specialization is shaped by experience, lateralization in higher-order motor areas should shift toward the right hemisphere in individuals with R-OBPI because of their reversed hand dominance. Critically, these effects should be specific to right-side injury but not general effects of OBPI, hence should not be observed in individuals with L-OBPI (**Fig. 1A**). We next considered three competing hypotheses regarding the consequences of motor reorganization for language lateralization. If action and language lateralization are constrained by common developmental mechanisms, language lateralization should shift toward the right hemisphere alongside motor lateralization in individuals with R-OBPI (**Fig. 1A**). Alternatively, if motor and language systems compete for neural resources, language lateralization should become more strongly left-lateralized as motor lateralization shifts rightward (Seydell-Greenwald et al., 2023; Cai et al., 2013; Lidzba et al., 2006). Finally, if the two systems emerge independently, language lateralization should remain unchanged.

## Results

Using functional MRI, we examined motor and language cerebral lateralization in individuals with OBPI and typical right-handed controls. Participants performed a manual action task that involved reaching-to-touch (reaching) or reaching-to-grasp (grasping) an object with their hands (**Fig. 1B**) and an auditory language task of sentence comprehension (**Fig. 1D**).

The critical group consisted of individuals with R-OBPI (**Fig. 1A**; N = 30, 15.0 ± 4.41 yrs), who developed lifelong left-hand dominance because of severe motor impairment of the right upper limb from birth. This unique natural experiment dissociates the brain’s presumed innate predisposition for left-hemisphere dominance from postnatal sensorimotor experience. A group of typical right-handers served as controls (**Fig. 1A**; N = 23, 25.3 ± 2.56 yrs).

As individuals with R-OBPI also exhibit an overall imbalanced hand use and impaired bimanual coordination, differences in cerebral lateralization may arise from these general deficits rather than postnatal left-hand dominance. To isolate these factors, a group of individuals with OBPI on the left upper limb (L-OBPI, **Fig. 1A**; N=22, 11.6 ± 3.27 yrs) was recruited. Unique hemispheric dominance patterns in the R-OBPI group relative to both controls and the L-OBPI group would support our core hypothesis that hemispheric specialization is shaped specifically by hand use experience. We first examined hemispheric lateralization during the action task, followed by language lateralization during sentence comprehension, and finally tested the relationship between motor and language lateralization across individuals.

### Motor experience alters the hemispheric dominance of frontoparietal motor areas

During the action task, participants reached toward and either touched or grasped a centrally located object with the left or right hand, depending on the instruction (**Fig. 1B**; **Methods**). Despite severe congenital motor impairment, most participants with OBPI were able to perform both actions with the affected hand and spent a longer time grasping versus simply touching the object (**Fig. 1C**, **SI Methods**), demonstrating successful differentiation between action types. We therefore focused on the grasping condition, which was expected to engage higher-order motor networks more strongly than simply reaching. Results from the reaching condition showed similar patterns and are presented in the **Supplementary Information (Fig. S1, S2, S4)**.

**Figure 2.**
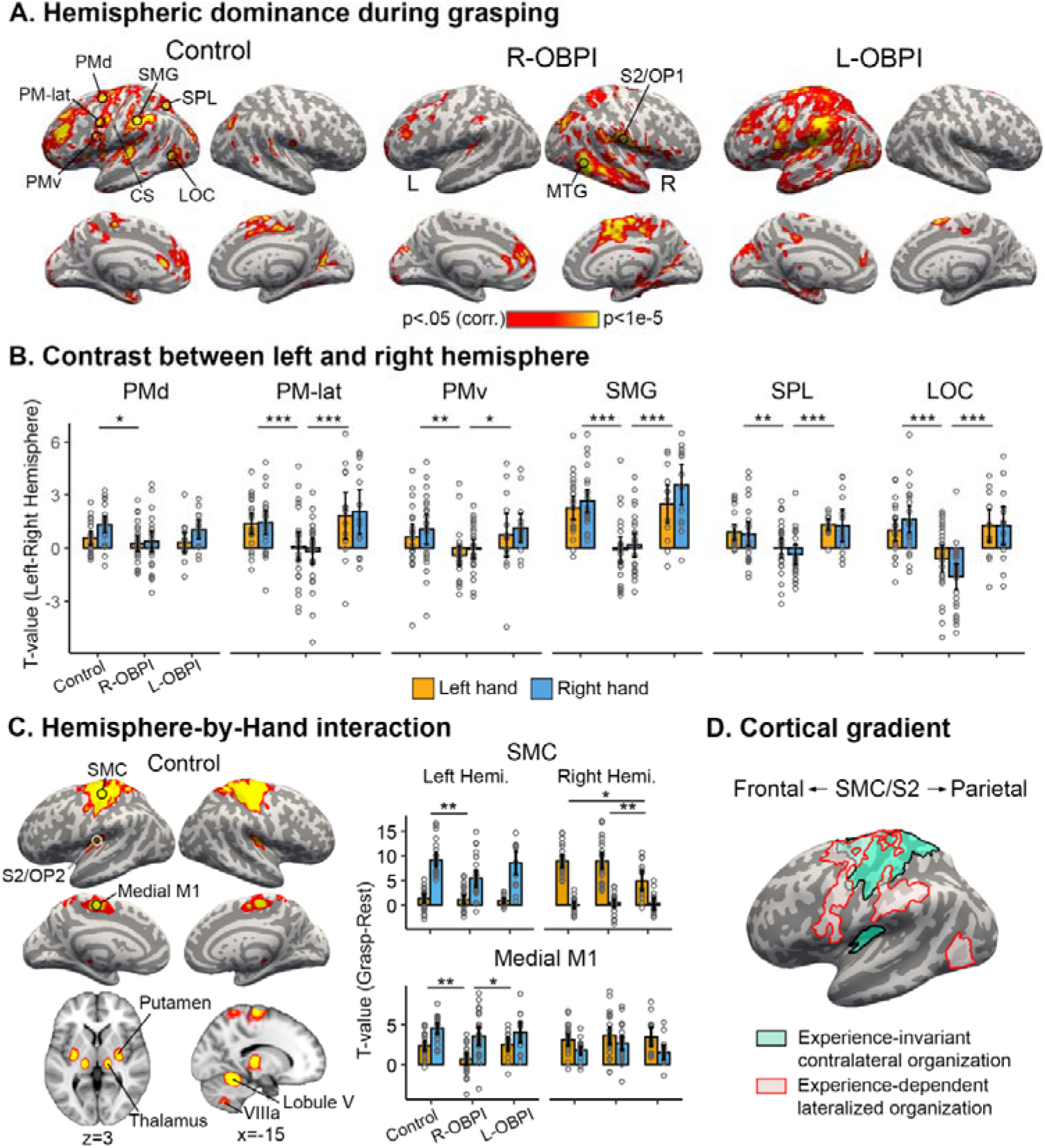
Experience-dependent and experience-invariant organization of hemispheric dominance in the motor system. **(A)** Hemispheric dominance maps during grasping. Controls showed pronounced left-hemisphere dominance across premotor, parietal, and lateral occipital cortices, while sparing the central sulcus (CS). This leftward dominance was substantially reduced in individuals with right-sided obstetric brachial plexus injury (R-OBPI), while preserved in individuals with left-sided OBPI (L-OBPI) Statistical maps show the contrast between original and flipped maps collapsed across hands (cluster-wise corrected at *p* < 0.05). **(B)** Mean hemispheric dominance values (left hemisphere minus right hemisphere) are shown for regions identified in controls, including dorsal premotor cortex (PMd), lateral premotor cortex (PM-lat), ventral premotor cortex (PMv), supramarginal gyrus (SMG), superior parietal lobule (SPL), and lateral occipital cortex (LOC). Positive values indicate stronger left-than right-hemisphere responses. Across ROIs, R-OBPI participants exhibited significantly reduced left-hemisphere dominance compared with both controls and L-OBPI participants, independent of the hand used for action. Each point is a participant. Error bars denote 95% CI. Circles indicate individual participants. Asterisks denote significant post hoc comparisons. * *p* < .05; ** *p* < .01; *** *p* < .001. Raw activation levels for each hand in each hemisphere are shown in Fig. S2. **(C)** Hemisphere-by-hand interaction reveals hand-dependent hemispheric dominance that is stable across groups (see **Fig. S3A** for maps from the OBPI groups). These regions, including primary sensorimotor cortex (SMC), secondary somatosensory cortex (S2), medial primary motor cortex (M1-medial), thalamus, and putamen, exhibited the expected contralateral organization (see **Fig. S3B** for ROI-level results in more ROIs). Cerebellar lobule V showed ipsilateral organization. Besides these experience-invariant patterns, ROI analyses of Hand × Group within each hemisphere additionally revealed reduced activation in the hemisphere contralateral to the injured limb, consistent with use-dependent plasticity. MNI coordinates are shown for maps displayed in the volumetric space. **(D)** Overlaying the main effect of Map and Hand × Map interaction from controls reveals a cortical gradient in the plasticity of motor networks. Whereas the canonical contralateral organization in SMC and secondary somatosensory cortex (S2/OP2) was stable across groups, the left-lateralized frontoparietal cortex and LOC in controls showed reorganization in face of altered hand use in R-OBPI.

We first quantified hemispheric lateralization by performing a voxel-wise mirror subtraction analysis (Baciu et al., 2005; Seghier & Price, 2011). Beta maps for the contrast of grasping versus rest were flipped along the left-right (X) axis and subtracted from the original maps. In the resulting difference maps, positive values in the left hemisphere indicate stronger activation in the left than the right hemisphere and positive values in the right hemisphere indicate right-hemisphere dominance. This approach therefore provides voxel-wise localization of hemispheric dominance.

For each group (Control, R-OBPI, L-OBPI), a random-effect ANOVA was performed with Map (original, flipped) and Hand (left, right) as fixed factors and participants as a random factor. A main effect of Map would identify regions exhibiting hand-independent hemispheric dominance, whereas a Map × Hand interaction would identify hand-dependent dominance, such as the canonical contralateral organization in primary sensorimotor cortex.

First, consistent with previous reports of a left-lateralized frontoparietal organization, Control group showed stronger activation in the left hemisphere in premotor, parietal and lateral occipital cortices, while sparing the primary somatosensory and motor cortex (**Fig. 2A**). Critically, this left lateralization was markedly reduced in the R-OBPI group but preserved in the L-OBPI group (**Fig. 2A**), suggesting specific impact of right-side OBPI. Corroborating these findings, across core sensorimotor regions-of-interest (ROIs) sampled from controls’ map, we found reduced left-hemisphere dominance in the R-OBPI group compared to both the Control and L-OBPI group independent of hand (**Fig. 2B**). R-OBPI group additionally showed strong dominance in right secondary primary somatosensory cortex (S2/operculum parietal OP1) and right middle temporal gyrus (MTG; **Fig. S3C**). Decomposition of responses by hemisphere revealed that the reduced left lateralization reflected mixed effects of decreased response in the left hemisphere and increased response in the right hemisphere (**Fig. S2**). The comparable hemispheric lateralization observed in the L-OBPI and control groups argues against explanations based on general motor impairments or age differences. These findings demonstrate that hemispheric lateralization in higher-order frontoparietal areas is shaped by postnatal sensorimotor experience, such that lifelong predominant use of the left hand reduced the characteristic left-hemisphere dominance and resulted in a more bilateral organization.

By contrast, the Map × Hand interaction revealed a network with hand-dependent lateralization that was preserved across all three groups (**Fig. 2C, Fig. S3A, S3B**). Specifically, the primary somatosensory and motor cortex (SMC), S2 (parietal operculum OP2), posterior putamen and thalamus showed typical contralateral representations while the cerebellum exhibited ipsilateral representations irrespective of group. Thus, unlike higher-order frontoparietal regions, hemispheric lateralization in primary sensorimotor and subcortical regions remained stable despite dramatically altered hand use experience (**Fig. 2D**). Nevertheless, these regions still showed local use-dependent plasticity, with a general reduction in activation level in the hemisphere contralateral to the injured limb (**Fig. 2C, Fig. S3B,** bar graph; **Methods)**, suggesting that experience modified regional responses without altering their canonical contralateral organization.

### Motor experience reorganizes bimanual action representations

We next quantified the spatial overlap between activations evoked by the two hands, hereafter referred to as bimanual recruitment. Previous studies have shown that higher-order frontoparietal regions encode action information in a hand-independent manner (Di Caro et al., 2025; Merrick et al., 2022; Kumar et al., 2020). We therefore asked whether the hemispheric dominance of bimanual recruitment is likewise shaped by postnatal hand use experience. Importantly, the hemispheric lateralization analysis above does not necessarily predict bimanual recruitment, because one hemisphere may exhibit stronger overall activation without engaging more overlap between the two hands.

First, for each group, we calculated an overlap probability map denoting the number of participants showing bimanual recruitment in each voxel. In both the Control and L-OBPI group, bimanual recruitment was concentrated in the left premotor, parietal, and lateral occipictal cortices (LOC), while largely sparing the central sulcus where SMC resides. In contrast, the R-OBPI group shows a more extensive and consistent pattern of right-hemisphere bimanual recruitment (**Fig. 3A, Fig. S4A**). To quantitatively compare the lateralization of bimanual recruitment, we sampled large-scale ROIs of premotor cortex, supplementary motor area (SMA), superior parietal lobe (SPL), inferior parietal lobe (IPL), and lateral occipical cortices in both hemispheres based on the Juelich Histological Atlas. For each ROI and group, we calculated a laterality index (LI) denoting which hemisphere contained more voxels exhibiting bimanual recruitment (**Methods**). Across all ROIs, the R-OBPI group showed negative LIs (significantly below zero in all but LOC), i.e. right-hemisphere dominance, relative to the Control and the L-OBPI group (**Fig. 3B, Fig. S4B**). Experience-dependent plasticity in SMA and SPL is further supported by even stronger left lateralization in the L-OBPI vs. controls, presumably as a consequence of enhanced right-hand use in L-OBPI. Taken together, these findings indicate that postnatal sensorimotor experience reshapes the hemispheric organization of hand-independent action representations rather than merely altering the magnitude of motor responses.

**Figure 3.**
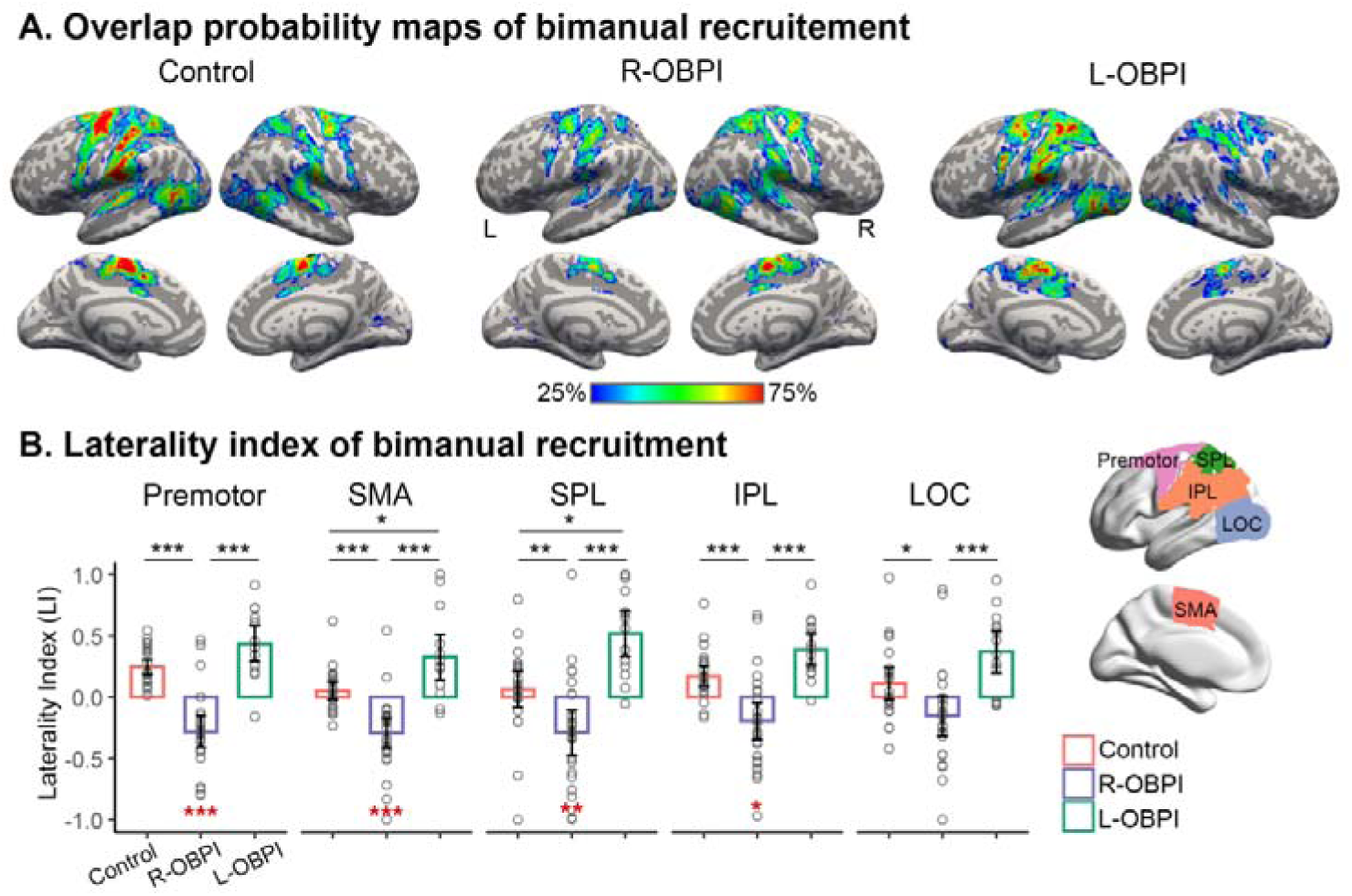
Experience-dependent reorganization of bimanual recruitment in higher-order motor regions. **(A)** Overlap probability maps of bimanual recruitment for each group. Maps indicate the proportion of participants showing significant activation in each voxel during movements of both hands. In Controls and the L-OBPI group, bimanual recruitment was concentrated in left premotor, parietal, and lateral occipital cortices. In contrast, the R-OBPI group exhibited a larger spatial extent and greater inter-individual overlap of bimanual recruitment in the right hemisphere. **(B)** Laterality indices (LI) of bimanual recruitment in bilateral premotor cortex, supplementary motor area (SMA), superior parietal lobule (SPL), inferior parietal lobule (IPL), and lateral occipital cortex (LOC). Positive values indicate left-hemisphere dominance. Relative to Controls and the L-OBPI group, the R-OBPI group showed significantly reduced or reversed lateralization. Each point is a participant. Red asterisks denote whether R-OBPI showed significantly negative LIs. Error bars denote 95% CI. * *p* < .05; ** *p* < .01; *** *p* < .001.

### Differentiation of action types in the unaffected hemisphere in the OBPI groups

We then investigated more fine-grained action representation by contrasting grasping and reaching actions across both hands, evaluated by a univariate analysis of Action (reaching, grasping) by Hand (left, right) ANOVA. In controls, a main effect of Action (driven by grasping > reaching) independent of hand was primarily found in a left-lateralized network encompassing the left superior parietal lobe and primary somatosensory cortex, along with bilateral medial SMC, S2, and cerebellum (**Fig. 4A**). In contrast, the corresponding action-selective responses were predominantly localized to the right SPL and SMC in individuals with R-OBPI (**Fig. 4A**). Although no main effect of Action was found in the L-OBPI group, within-hand contrasts consistently localized to the left S1 for both hands, showing preserved left-lateralization (**Fig. 4A**).

**Figure 4.**
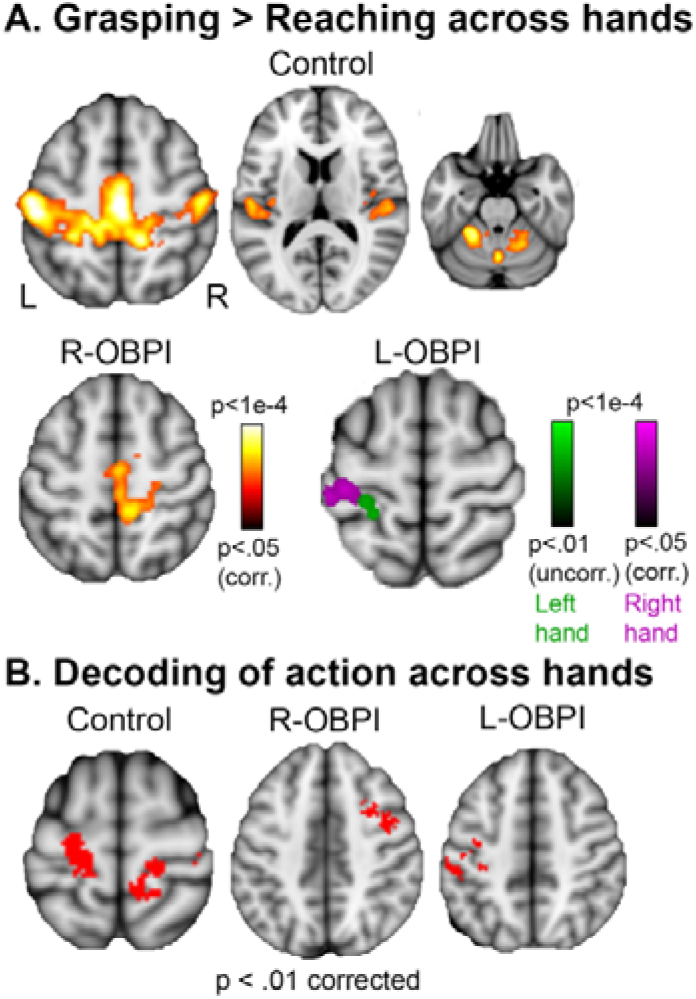
Differentiation between grasping and reaching in each group. **(A)** In controls, grasping elicited stronger activation in left in bilateral SMC, medial SMC, S1, S2, and cerebellum compared with reaching. These effects shifted toward the hemisphere contralateral to the dominant hand in both OBPI groups, i.e. in right SPL and S1 in the R-OBPI group and in left SPL and S1 in L-OBPI. **(B)** Whole-brain searchlight decoding of action type (reaching, grasping) across hand in each group. In controls, hand-independent action-type representation was found in a bilateral network, whereas in the OBPI group these representations were primarily found in the hemisphere contralateral to the dominant hand, i.e. in right PMd in the R-OBPI group and in left intraparietal sulcus and SMC in the L-OBPI group.

To directly probe hand-independent representational content of each action, we performed cross-hand multivoxel pattern analysis (MVPA), training classifiers between reaching and grasping with one hand and testing on the other hand (**Methods**). Whereas controls demonstrated cross-hand action decoding in bilateral sensorimotor areas, the OBPI groups primarily showed hand-independent action representation in the hemisphere contralateral to the dominant hand (i.e. right hemisphere in R-OBPI and vice versa; **Fig. 4B**). Together, these findings demonstrate that developmental hand use experience reorganizes abstract, hand-independent action representations rather than merely altering motor activation patterns.

### Reduced left-hemisphere lateralization of language in BA44 in R-OBPI

Does the experience-dependent reorganization observed in the motor system extend to the language network? We next evaluated brain responses from the fMRI language experiment to address this question. Participants listened to sentences or backward speech and performed a judgment task after each sentence (**Fig. 1D**; **Methods**). The sentences consisted of simple, age-appropriate descriptions of third-person events without reference to the participants’ own actions, with minimal semantic overlap with the action task (**SI Methods**). Brain response during sentence listening and backward speech was contrasted to localize the language network. All three groups showed language selectivity (i.e. sentence vs. backward speech) in a typical left-lateralized frontotemporal network including middle and inferior frontal gyrus, and middle and posterior temporal gyrus (**Fig. S5**), consistent with past findings (Malik-Moraleda et al., 2022; Lipkin et al., 2022; Olulade et al., 2020).

We quantified voxel-wise language lateralization using the voxel-wise mirror subtraction analysis and compared language lateralization across groups (**Fig. 5A**). If the effect of postnatal hand use experience propagates to the language system, reduced left lateralization in the frontotemporal language network would be observed in the R-OBPI group relative to the Control and L-OBPI group. Strikingly, at both whole-brain and ROI level, R-OBPI showed reduced left-hemisphere dominance in left IFG pars opercularis (BA44, Broca’s area) relative to both controls and L-OBPI (**Fig. 5B**). There were also significantly more individuals with R-OBPI that showed right-hemisphere dominance (44.8%) relative to controls and L-OBPI (0% and 4.76% respectively, Fisher–Freeman–Halton exact test: *p* < .001; see **Fig. S7-S9** for individual activation maps of each group).

**Figure 5.**
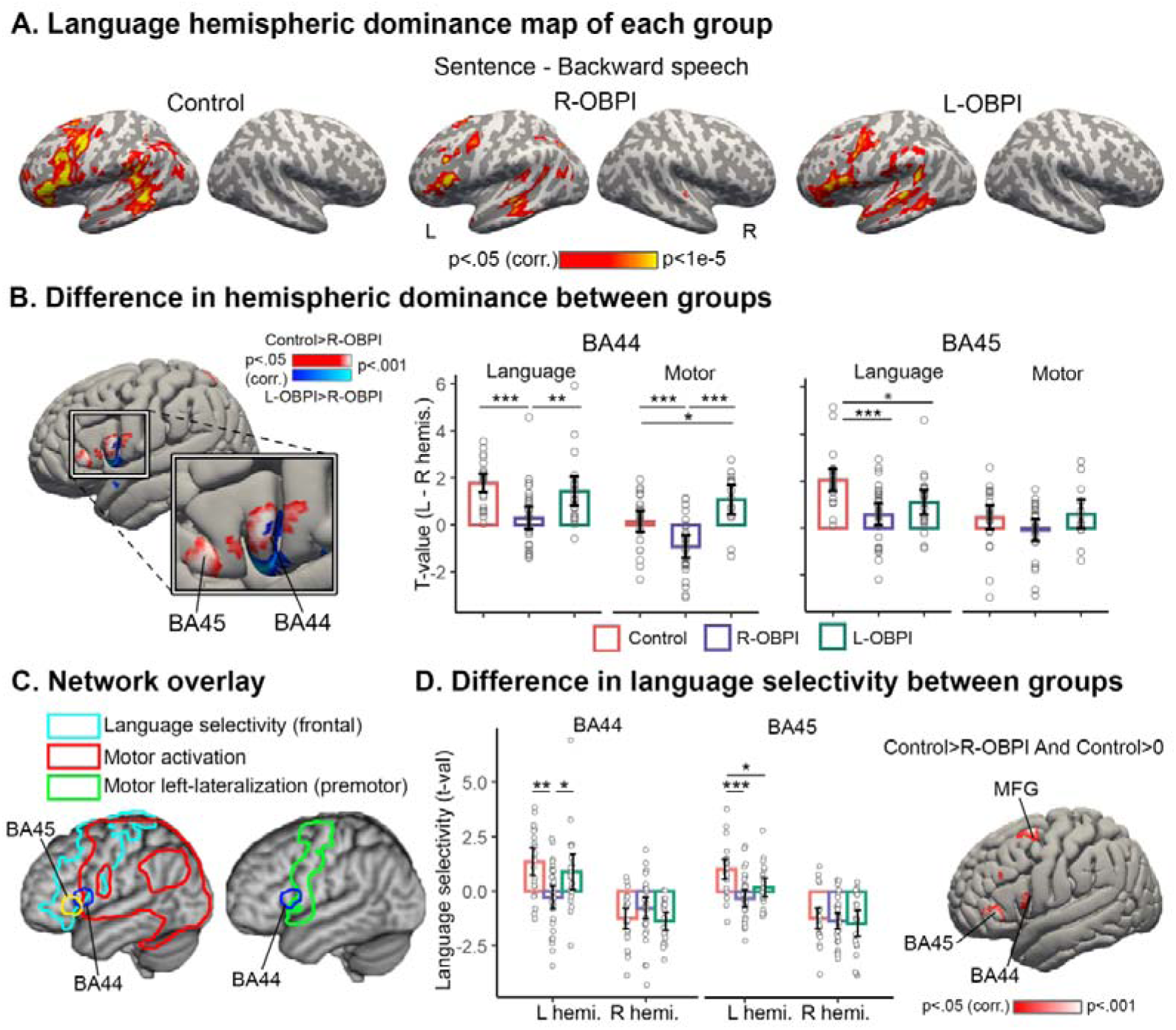
Reduced left-hemisphere lateralization and language selectivity of language processing in R-OBPI. **(A)** Hemispheric dominance map for each group. All groups exhibited left-lateralization within a predominantly left-lateralized frontotemporal network encompassing inferior and middle frontal gyri and middle/posterior temporal cortices. **(B)** Whole-brain voxel-wise comparisons revealed reduced left-hemisphere dominance in R-OBPI relative to both Controls and L-OBPI in BA44 (Broca’s area), along with reduced left-lateralization for grasping. In BA45, both OBPI showed decreased left-lateralization for language compared with controls, with no difference in motor lateralization across groups. Each point is a participant. Error bars denote 95% CI. * *p* < .05; ** *p* < .01; *** *p* < .001. **(C)** BA44 is located at the interface between the frontal language-selective network (outlined in cyan, defined as controls’ language selectivity map, **Fig. S5A**) and the motor network (outlined in red, defined as the right-hand grasping activation map from controls), whereas BA45 belonged exclusively to the language network. In addition, BA44 is located just anterior to the frontal left-lateralized motor network (outlined in green, based on Fig. 2B), showing no hemispheric lateralization despite motor activation. **(D)** Whole-brain analysis identified reduced language selectivity in R-OBPI relative to Controls within left BA44, BA45, and middle frontal gyrus (MFG). ROI analyses revealted that the reduced lateralization was primarily driven by decreased language selectivity in the left hemisphere rather than increased right-hemisphere responses. Unlike BA44, BA45 and MFG exhibited reduced language selectivity in both OBPI groups compared with Control (**Fig. S10B**).

The anatomical location and functional profile of BA44 make it a region of interest for understanding variability in hemispheric lateralization across individuals. In controls, this area intersects the motor and language network, being active in both tasks (**Fig. 5C, Fig. S10A)**. Notably, it locates just anterior to the left-lateralized frontal motor network shown in **Fig. 2A**, showing strong left-lateralization in language but not in action (**Fig. 5B, 5C**). Nevertheless, its action dominance shifted toward right in R-OBPI and toward left in L-OBPI (**Fig.5B**), mirroring the reorganization patterns with other frontoparietal motor areas (**Fig. 2A and 2B**). These results raise the possibility that IFG BA44 is potentially an interface area between higher-order action and language system in which reorganization driven by altered motor experience propagates to the language system.

BA44 has been implicated in speech production (Hickok & Poeppel, 2004; Fedorenko et al., 2024), raising the possibility that the neural responses observed during sentence listening may partly reflect implicit articulatory processes rather than language processing per se. First, our BA44 is located anterior to the speech-production network based on the Neurosynth meta-analytic database (https://neurosynth.org/; Yarkoni et al., 2011), intersecting the speech motor system and higher-order language parcels defined from large-sample functional localizer studies (**Fig. S6A**; Fedorenko et al., 2010). Importantly, the reduced left lateralization in R-OBPI was replicated when BA44 was restricted to its intersection with the independently defined IFG language parcel (**Fig. S6A, BA44 (IFG)**; Fedorenko et al., 2010), demonstrating that the effect remained robust after excluding adjacent motor-related regions. Second, we constructed representational dissimilariy matrices (RDMs) across sentence stimuli based on their articulatory dynamics (**Fig. S6B**; **SI Methods**) and found no evidence for correlation with the neural RDMs in BA44 (Control: Pearson’s *r* = 0.003, permutation *p* = .339; R-OBPI: *r* = 0.008, *p* = .121; L-OBPI: *r* = -0.004, *p* = .681). Together, these findings suggest that the BA44 effect reflects language-related processing that is not readily explained by articulatory or motor representations alone.

We next tested whether reduced left lateralization reflected weaker left-hemisphere responses or stronger right-hemisphere responses. ROI analyses in homologous areas of IFG BA44 revealed that the effects were primarily driven by decreased language selectivity in the left hemisphere (**Fig. 5D**, bar graph). These results were further confirmed in whole-brain analyses comparing language selectivity across groups (**Fig. 5D**, brain map).

In addition to BA44, BA45 also exhibited reduced left lateralization in R-OBPI groups relative to controls (**Fig. 5B**), an area that belongs exclusively to the language network and showed no motor activation (**Fig. 5C, Fig. S10A**). Unlike BA44, however, BA45 showed no evidence of differential lateralization between the two OBPI groups (Post-hoc Tukey-adjusted comparison of R-OBPI vs. L-OBPI: *t*(70) = -1.50, *p* = .300; **Fig. 5B**). Rather, both OBPI groups showed reduced lateralization and language selectivity in BA45 than controls (**Fig. 5B**, **Fig. 5D, bar graph**), suggesting that reduced lateralization in BA45 may reflect a more general characteristic of the OBPI population.

Finally, we explored hemispheric lateralization in independent, commonly used pre-defined language parcels (Fedorenko et al., 2010; **Fig. S6A**) and in basal ganglia that showed overlapping activation of tool-use and syntax processing (Thibault et al., 2021). In both IFG and IFGorb, individuals with R-OBPI showed significantly reduced left lateralization relative to controls, with a similar albeit non-significant trend relative to L-OBPI. These results therefore potentially extend findings on BA44 to a broader inferior frontal language network. No group differences were found in basal ganglia (**Fig. S6A**).

Despite differences at the neural level, no differences in language performance were detected between the R-OBPI and L-OBPI groups in either sentence judgment accuracy or reaction time (**Fig. 1E**), including analyses restricted to age-matched participants (**Fig. S11A, C**). Both OBPI groups performed less accurately and more slowly than the Control group (**Fig. 1E**), likely reflecting their younger age (**Fig. 1A**).

### Correlation in left-hemispheric dominance between language and action

Finally, we examined whether individual differences in language lateralization were associated with lateralization of the motor system. If the reduced left-lateralization of language in R-OBPI reflects shared developmental constraints with the action system, individuals with stronger left-lateralization of action should also exhibit stronger left-lateralization of language. Alternatively, if altered hand use merely increases variability in hemispheric organization (Labache et al., 2020), no systematic relationship would be expected.

We focused on language lateralization in BA44 and correlated it with action lateralization in BA44 itself and in frontoparietal motor ROIs showing a main effect of Map in the Hand × Map ANOVA (**Fig. 2B).** Consistent with our hypothesis, language and motor lateralization were positively coupled specifically in individuals with R-OBPI, in left PMd (partial correlation controlling for age, Pearson’s *r* = 0.43, *p =* .04) and lateral PMd (Pearson’s *r* = 0.52, *p =* .01). This action lateralization was quantified across the two hands, but the affected and unaffected hand may undergo distinct reorganization patterns. We therefore examined left- and right-hand motor lateralization separately. Strikingly, language lateralization was selectively associated with left-hand action lateralization across several frontal motor regions, including BA44 and dorsal, lateral, and ventral premotor cortex, with similar but weaker trends observed in parietal cortex (**Fig. 6A and 6B; Fig. S11A**). In contrast, no correlation was observed for the right-hand despite comparable activation level with the left hand (**Fig. S2**, **Fig. S12A**). Moreover, these positive correlations were specific to the R-OBPI group, whereas controls and L-OBPI showed weak or negative relationships (**Fig. 6A, Fig. S12A**). Finally, these correlations were limited to BA44 and were not consistently observed in BA45 (**Fig. S12B**).

**Figure 6.**
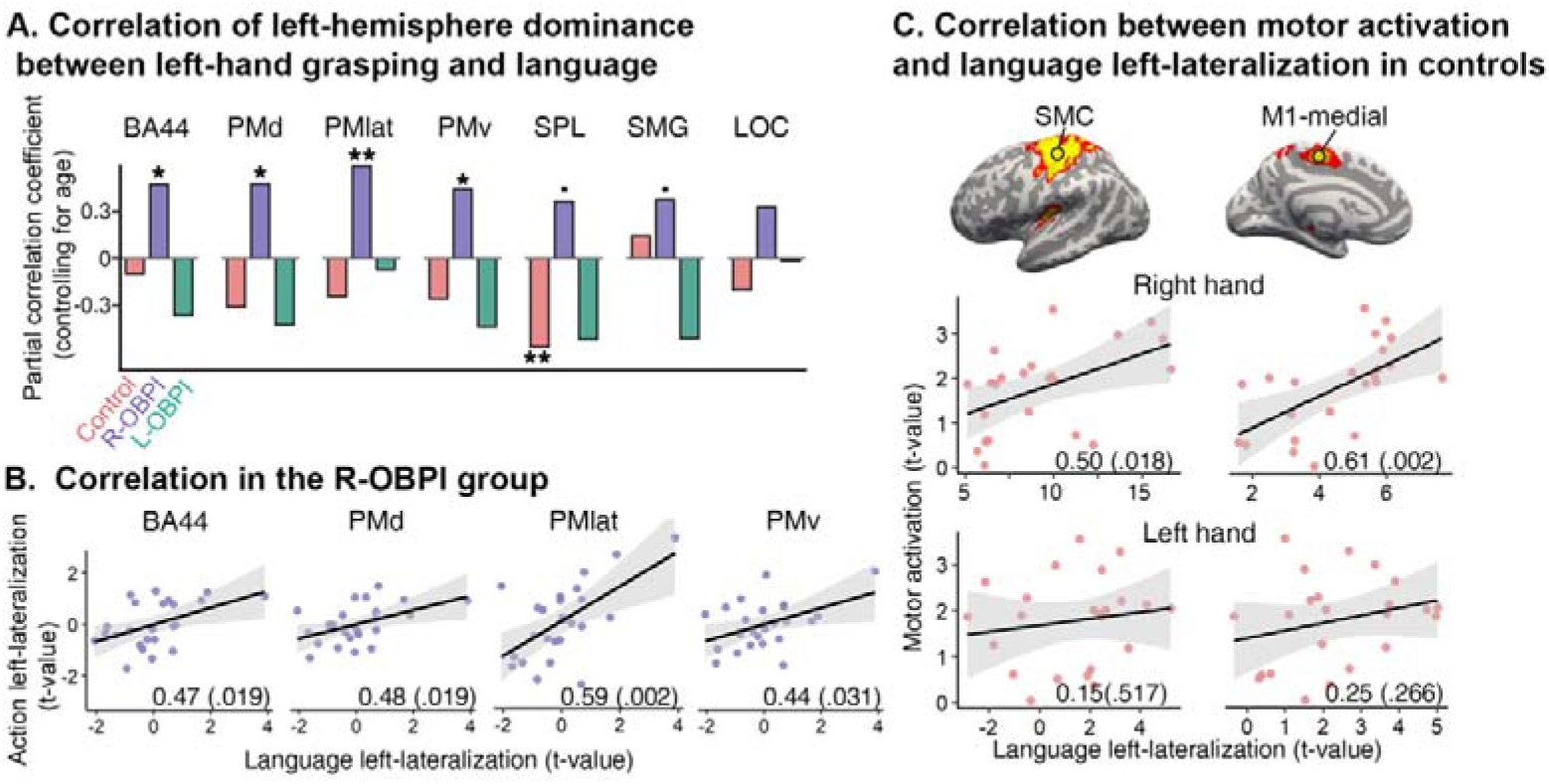
Correlation between language left-lateralization in BA44 and motor left-lateralization across motor ROIs. **(A)** Partial correlation (controlling for age) between language left-lateralization in BA44 and motor left-lateralization (for the left hand) in BA44 as well as higher-order motor areas (same ROIs as from Fig. 2B). In R-OBPI, language lateralization was positively associated with left-hand motor lateralization across frontal motor regions including BA44, PMd, PMlat, and PMv, and the same trend in SPL and SMG. \**p* < .05, \*\**p* < .01, \*\*\**p* < .001, ·*p* < .1. See **Fig. S11** for correlation with right-hand and hand-averaged motor lateralization. **(B)** Scatter plots of significant correlations in (A), between language and left-hand motor lateralization. Each point is a participant. Shaded areas denote 95% CI. Numbers at the bottom represent Pearson’s correlation coefficients (*r*), with the corresponding *p*-values shown in parentheses. Scatter plots show residual correlation after controlling for age. **(C)** Controls showed a positive association between language lateralization in left BA44 and motor responses to right-hand grasping in left SMC (left column) and medial M1 (right column). Each point represents a participant. Shaded areas denote 95% CI.

Besides the frontoparietal areas, local use-dependent plasticity occurred in R-OBPI in left SMC, medial M1, S2, and cerebellum, with a general reduction in motor activation compared with controls and L-OBPI (**Fig. 2C**). We then explored whether language lateralization was associated with activity changes in these lower-level motor regions. Surprisingly, language lateralization in left BA44 and BA45 was associated with motor responses in left SMC and medial M1, particularly for the right hand movements and in controls (**Fig. 6C, Fig. S12C**). No correlation was found for other groups (**Fig. S12D**). These exploratory findings converge on a common relationship between language lateralization and the organization of motor circuits supporting the functional hand. In R-OBPI, this relationship was observed in higher-order motor regions for the compensatory left hand, whereas in controls it was observed in lower-level motor regions for the dominant right hand.

Together, these findings demonstrate that experience-dependent reorganization is coordinated across motor and language systems at the individual level, providing direct evidence that hemispheric specialization across cognitive domains is shaped by common developmental mechanisms.

## Discussion

By leveraging a unique natural experiment in which innate predisposition for hemispheric dominance is dissociated from hand-use experience, we demonstrated that postnatal asymmetric sensorimotor experience causally reshapes cerebral lateralization. This reorganization was selective, affecting higher-order frontoparietal motor areas while sparing the primary sensorimotor and subcortical regions. Across multiple analyses, including hemispheric response strength, bimanual recruitment, and univariate and multivariate action type differentiation, individuals with R-OBPI consistently showed reduced or reversed left-hemisphere dominance relative to controls and individuals with L-OBPI. Critically, experience-dependent reorganization extended to the language network, resulting in reduced left lateralization of BA44 during sentence comprehension specifically in individuals with R-OBPI. Moreover, language lateralization in BA44 was correlated with motor lateralization across frontoparietal motor areas in individuals with R-OBPI. Taken together, these findings demonstrate that hemispheric specialization is shaped by sensorimotor experience during development and provide evidence that experience-driven reorganization can propagate across cognitive domains, supporting coordinated rather than independent development of action and language systems.

### Hierarchical principles of experience-driven motor plasticity

Higher-order motor regions have long been implicated in the left-hemisphere specialization of human action (Di Caro et al., 2025; Merrick et al., 2022; Buxbaum & Saffran, 2002; Johnson-Frey 2004). A critical question is how left-hemisphere dominance emerges during development. Although both innate predispositions and developmental experience have long been proposed to contribute to hemispheric specialization (Corballis, 2014; Ocklenburg et al., 2014), their relative contributions have remained difficult to disentangle because genetic, prenatal, and postnatal influences are typically intertwined. By exploiting unilateral OBPI as a natural experiment that selectively alters postnatal hand use while preserving the genetic predisposition, we showed that individuals with R-OBPI showed markedly reduced left-hemisphere dominance compared with controls and individuals with L-OBPI. However, this reduction resulted in a more bilateral pattern rather than a reversal toward right lateralization, suggesting preserved intrinsic contraints of left-hemisphere dominance. These findings suggest that sensorimotor experience shapes hemispheric specialization without overriding the intrinsic bias of left-hemisphere dominance.

One possible developmental account is that higher-order action representations are established bilaterally early in development but with an initial advantage in the left hemisphere (Kosslyn, 1987). During typical development, this initial left-hemisphere advantage and preferential use of the right hand reciprocally reinforce each other, presumably through recurrent interactions between higher-order motor regions and sensorimotor circuits. In contrast, lifelong reliance on the left hand in individuals with R-OBPI may alter this developmental trajectory through a combination of reduced specialization of the left hemisphere and increased engagement of homologous regions in the right hemisphere. The combination of these changes weakens the original leftward bias and produces a more bilateral organization. Developmental neuroimaging studies similarly suggest that higher-order action networks become increasingly left-lateralized from childhood to adulthood (Biagi et al., 2015; also see Olulade et al., 2020). Our findings further suggest that this developmental transition is actively shaped by sensorimotor experience rather than reflecting passive maturation alone. More broadly, these findings are consistent with the view that cortical organization is largely constrained by innate connections but subsequent relevant experience is required to maintain and refine such organization (Striem-Amit et al., 2018; Dall’Orso et al., 2018; Arcaro et al., 2017a; 2017b; 2019). As our participants with OBPI involves both early complete paralysis and subsequent compensatory use of the unaffected hand, future investigations of individuals with milder OBPI who recover near-normal function within the first months of life will help distinguish the effect of early sensorimotor deprivation versus later hand use in the organization of motor networks (Tucciarelli et al., 2025).

Higher-order frontoparietal motor areas are considered relatively independent of specific sensorimotor experience, encoding function-based and effector-independent abstract action representations. In individuals born without both hands and use their feet for daily actions, the frontoparietal areas represent actions performed with the feet in manners similar to hand actions in typical controls (Liu et al., 2020; Martinez-Addiego et al., 2025). At first glance, the present findings appear to contrast with previous studies. If higher-order motor areas developed independently of experience, altered hand use in R-OBPI would not be expected to affect hemispheric specialization. We propose that these findings can be reconciled by considering the different aspects of cortical organization they address. Previous studies manipualted the acting effector while keeping hemispheric dominance effectively constant, examining the left hemisphere in response to movement performed by the right side of the body (Martinez-Addiego et al., 2025; Liu et al., 2020; Gallivan et al., 2013). In contrast, the present study manipulated the balance of sensorimotor experience between the two hemispheres and examined hemispheric organization. Consistent with previous work, our MVPA results demonstrate hand-independent action representations in higher-order motor cortex in the dominant hemisphere in each group. At the same time, we show that the hemispheric implementation of these computations remains developmentally plastic. Therefore, postnatal experience appears to influence not whether higher-order action representations emerge, but which hemisphere preferentially supports them.

In contrast to the extensive reorganization observed in higher-order motor cortex, primary sensorimotor cortex retained its canonical contralateral organization across all groups. Previous studies suggest that the primary sensorimotor hand area is already established by birth (Dall’Orso et al., 2018; Arcaro et al., 2019). In individuals born without a hand, however, this territory reorganizes to represent other body parts (Striem-Amit et al., 2018; Hahamy et al., 2017), indicating that developmental sensorimotor experience is required to maintain hand selectivity (Tucciarelli et al., 2025). Once established, the hand representation remains remarkably stable in acquired amputees despite decades without peripheral input (Shone & Makin, 2025; Kikkert et al., 2016). In our participants with OBPI, the affected hand received little or no voluntary sensorimotor input during the first months of life and only partially recovered basic movements following surgery. Despite this early-life paralysis and persistent limitations in hand function, contralateral hand representations were preserved, extending previous findings that primary sensorimotor hand representations remain relatively stable following major alterations in sensorimotor experience.

### Coupled organization of action and language lateralization

Beyond its effects on the motor system, early sensorimotor experience also shapes the developmental organization of the language network. Language lateralization develops gradually throughout childhood, with robust left lateralization emerging during the first few years of life and continuing to mature throughout childhood (Berl et al., 2014; Olulade et al., 2020; Ozernov-Palchik et al., 2026; Dehaene-Lambertz et al., 2006). This prolonged developmental trajectory provides a period during which experience can shape the functional organization of language networks. Consistent with this view, reduced functional selectivity has been reported in left-lateralized language regions following atypical early language experience in congenitally deaf individuals (Wang et al., 2023), and altered regional lateralization has been observed across the language network in early deafness (Yang et al., 2024). Our findings extend this developmental perspective by showing that altered sensorimotor experience can also modify the hemispheric organization of BA44, despite typical language exposure and a predisposition for left-hemisphere lateralization.

The relationship between language and action lateralization offers a mechanistic interpretation of how altered sensorimotor experience may influence language organization. In individuals with R-OBPI, language lateralization in left BA44 was positively associated with motor left lateralization, particularly in frontal motor areas and for movements of the unaffected left hand. Importantly, this association was not driven by motor activation per se, as both hands elicited comparable activation in frontal motor regions (**Fig. S2**). Rather, it may reflect the preservation of experience-dependent, functional motor representations in the left frontal cortex driven by the dominant use of the unaffected left hand in cases of impaired right-hand motor function in R-OBPI. This interpretation is consistent with views of functional links between language and action, such as hierarchical sequencing (Greenfield, 1991; Thibault et al., 2021; Friederici, 2023) and goal-directed feedback control (Hickok, 2012). Therefore, it may be the functional representations maintained in frontal motor areas, shaped by compensatory reliance on the functionally dominant hand, that are critical for their developmental coupling with the language system. This perspective suggests that the body serves not only as an interface for interacting with the world but also as a developmental scaffold for abstract cognitive systems such as language. Future studies employing a broader range of motor and language tasks that systematically vary their sensorimotor, semantic, and syntactic demands will help disentangle which functional components of action and language are linked through shared developmental mechanisms, and whether similar experience-dependent reorganization extends to other language domains or even broader cognitive systems (Fedorenko et al., 2020; 2024).

The findings in controls further suggest the importance of behavioral relevance of motor representations in shaping the relationship between action and language systems. In controls, language lateralization in BA44 was associated with responses to the dominant right hand, specifically in the contralateral left primary sensorimotor cortex where the dominant-hand representation is typically localized. Together with the findings in R-OBPI, these results suggest that language organization is linked to the motor representation that is most functionally relevant for everyday behavior, regardless of which hand supports this function. However, the cortical locus of this relationship differed across developmental conditions. Whereas this association involved the primary sensorimotor cortex in controls, it involved frontal motor areas involved in higher-order action planning in individuals with R-OBPI. Although the basis of this distinction remains unclear, several, not mutually exclusive, mechanisms may account for it.First, altered hand use in individuals with R-OBPI may increase developmental variation in higher-order motor organization, unmasking its covariance with that of the language network. Second, primary sensorimotor cortex may be relatively constrained by its predominantly contralateral corticospinal organization, whereas higher-order motor regions retain greater capacity for experience-dependent reorganization. Consequently, long-term adaptation to altered hand use may be expressed more prominently in premotor cortex, shifting the motor correlate of language lateralization from primary sensorimotor to higher-order motor regions.

The pattern observed in BA45 further suggests a gradient in how sensorimotor experience influences different components of the language network. Unlike BA44, BA45, located further anterior to the motor cortex and showed no motor activity (Pritchett et al., 2018), showed reduced left lateralization and language selectivity in both OBPI groups. This pattern suggests that regions less directly associated with action-related computations may be influenced by more general consequences of atypical sensorimotor development rather than experience-specific reorganization related to hand use. Such effects may reflect altered opportunities for bimanual coordination and complex object interaction following brachial plexus injury. Alternatively, they may partly reflect the younger age of the OBPI groups relative to controls and a less mature language network. Although language lateralization continues to develop throughout childhood (Berl et al., 2014; Ozernov-Palchik et al., 2026), current developmental evidence suggests that maturation is driven primarily by reductions in right-hemisphere language responses rather than increases in left-hemisphere activation (Olulade et al., 2020), and the overall lateralization remains stable since age four (Ozernov-Palchik et al., 2026), making age alone an incomplete explanation. Future studies with age-matched controls will be required to distinguish these possibilities.

### Broader implications for development and embodied cognition

Our findings have broader implications for understanding how early sensorimotor experience contributes to cognitive development. A large body of developmental research has demonstrated that early motor abilities predict later language outcomes (e.g., Iverson, 2010; Walle & Campos, 2014; Contino et al., 2025; Tseng & Hsu, 2025), yet the mechanisms underlying this relationship remain debated. One account emphasizes developmental cascades, whereby emerging motor skills enrich interactions with caregivers and increase opportunities for language learning (Iverson, 2010; Masten & Cicchetti, 2010; Libertus & Violi, 2016). Another proposes that action and language rely on shared neural computations, including hierarchical sequencing, structured action planning, gesture, and tool use (Citro et al., 2026; Friederici, 2023; Thibault et al., 2021; Greenfield, 1991). Our findings provide neural evidence supporting a complementary mechanism: developmental sensorimotor experience shapes the functional organization of higher-order frontal motor circuits, which in turn is associated with the hemispheric specialization of the language network.

More broadly, these findings extend theories of embodied cognition from online sensorimotor grounding to long-term brain development (Wilson, 2002; Allen et al., 2024). Rather than merely influencing cognition during task performance, sensorimotor experience appears to contribute to the developmental establishment of higher-order cortical organization while operating under intrinsic organizational constraints. This perspective suggests that bodily experience contributes to cognitive development by shaping the organizational principles through which abstract functions emerge. Understanding how early physical experience organizes the developing brain may therefore provide a unifying framework linking developmental neuroscience, embodied cognition, and the emerging field of embodied artificial intelligence.

## Methods

### Participants

All participants had unilateral OBPI and underwent individualized surgery — including nerve grafting, nerve transfer, or neurolysis (**SI Appendix A**) — at the Department of Hand Surgery at Huashan Hospital during infancy. They had a postoperative follow-up of at least 7 years and were aged ≥8 years at the time of testing. Although motor recovery of the affected limb remained incomplete, with persistent impairments in upper limb function at study entry, residual motor function was sufficient for these patients to complete the required fMRI tasks. Exclusion criteria were concurrent or prior neurological disorders (e.g., central palsy, cerebral hemorrhage, hydrocephalus) confirmed by definitive neurological reports and EEG, CT, or MRI findings, and participation in another clinical trial within the previous month.

Thirty individuals with right OBPI participated in the study (15.0 ± 4.41 years, 11 females). For the action task, one participant did not finish the experiment due to discomfort in the scanner. Four participants were excluded due to excessive head motion in both runs, resulting in 25 participants in the final analysis. For the language task, one participant was excluded from the language task due to an overall high miss rate (50%) and below-chance accuracy (33%) for the non-missed trials, resulting in 29 participants in the final analysis.

Twenty-two L-OBPI participated in the study (11.6 ± 3.27 years, 6 females). For the action task, seven participants were excluded from due to excessive head motion in both runs. One participant declined to perform the task, resulting in 14 participants in the final analysis. For the language task, one participant did not complete the experiment due to discomfort, resulting in 21 participants in the final analysis. Twenty-three typical controls participated in the study (25.3 ± 2.56 years, 13 females), all right-handed as measured with the Edinburgh Inventory (scores >40).

The three groups differ in age, with controls the oldest and individuals with L-OBPI the youngest (**Fig. 1A**, main effect of group: *F* (2,72) = 90.0, *p* < .001; post-hoc pairwise comparison *p*s < .003). Importantly, our conclusions are not confounded by age differences because we focused on the difference between the R-OBPI group relative to both the Control and the L-OBPI group.

All studies were approved by the Institutional Review Board of the Institute of Neuroscience, Chinese Academy of Sciences and of the Huashan Hospital. Written informed consent was obtained from all adult participants and from the parents or legal guardians of minor participants prior to participation.

### Experimental Design and Procedures

The action task was a 2 (Hand: left, right) x 3 (Action: reaching, grasping, writing) factorial design. The current study focusing only on reaching and grasping. A ball (5cm diameter) mounted on a rod was placed inside the scanner above the participant’s chest served as the target object (**Fig. 1B**). When possible, the participant’s head was tilted up a small degree to see the ball. A slow event-related design was adopted. At the beginning of each trial, the participant was played an auditory cue (“left” or “right”) informing them the hand to use and meanwhile shown an image depicting reaching or grasping on the screen for 800ms (**Fig. 1B**). Participants reached toward and touched the ball with the fingertips for reaching and grasped the ball into the palm for grasping. The image was presented to the left or right of the central square fixation depending on the hand, consistent with the auditory cue. The participant had 3 seconds to execute the action from the onset of the auditory and visual instructions, and remained at the end posture until hearing a beep (200ms) when they returned the hand to the home position on the belly. Inter-trial interval was randomly selected from 6s, 6.5s, and 7s. Each of the six conditions was repeated five times within each run in randomized orders, and each participant performed two runs. Each run was video-recorded at 30fps using an MRI-compatible CCTV camera (Shenzhen Sinorad Medical Electronic Co., Ltd.) installed inside the MRI room for offline analyses of movement parameters (**SI Methods**).

The language task was 2 (Syntactic difficulty: canonical, noncanonical) and 2 (Semantic plausibility: plausible, implausible) design along with a backward speech condition (**SI Methods**; Skeide et al., 2014; Thibault et al., 2021). A slow event-related design was used. In each trial, participants listened to either a sentence or backward speech about 3 seconds long (e.g. “A fox chased the chicken into a hole”). Five hundred milliseconds following each sentence, a short probe sentence was played via headphones as well as shown on the screen to the participant (e.g. “The fox ran into a hole”) and the participant indicated if the two sentences were congruent by pressing one of two buttons. The hand used for making responses was counterbalanced across controls, and individuals with OBPI pressed the buttons with the unaffected hand. The offset of a sentence and the onset of the next trial was spaced by 9 seconds, and participants were given a maximum of 4.5 seconds to respond. In trials with backward speech, speech was played within 3 seconds and inter-trial intervals were 4.5 seconds. Each of the five conditions was repeated five times within each run in randomized orders, and each participant performed two runs.

### MRI acquisition

Imaging data were collected in a Siemens Prisma 3T scanner at the Institute of Neuroscience, Chinese Academy of Sciences using a 32-channel head coil. T1-weighted structural images were collected using an MPRAGE sequence (voxel size: 1mm isotropic, TR = 2300ms, TE = 2.98ms, Flip angle = 9 deg). Functional images were collected using an echo-planar imaging sequence (voxel size: 2x2x2mm^3^, TR = 1500ms, TE = 30ms, Flip angle = 66 deg, no. of slices = 54).

## Data analysis

### Behavioral analyses

#### Action task

Participants’ motor performance was analyzed based on videos recorded during the experiment (**SI Methods**). Movement duration was calculated as the duration between movement onset and movement offset. A linear mixed-effects model analysis on trial-wise movement duration was performed with Action and Hand as within-subject factors, Group (Control, R-OBPI, L-OBPI) as a between-subject factor, and individual participants as random factors. Post-hoc analyses were then performed to examine the effect of Action and Hand within each group (**Fig. 1C**). All statistical analyses were ran in Rstudio (2021.09.0).

#### Language task

Accuracy and log-transformed reaction time were analyzed using linear mixed-effects models with Group as a between-subject factor and individual participants as random factors (**Fig. 1E, Fig. S12**). Trials with no response were excluded for the analysis of accuracy, and only correct trials were considered for the analysis of reaction time. In a second analysis, linear-mixed models were performed on accuracy and reaction time with Syntax (canonical, noncanonical) and Semantics (plausible, implausible) as fixed factors and participants as a random factor (**SI Methods; Fig. S12**).

### fMRI preprocessing

Neuroimaging data were preprocessed using FSL 6.0.5. Structural images were brain-extracted and co-registered to the MNI brain template (1mm isotropic) using 12-DOF linear transformation. Tissue-segmentation was performed using FAST. Functional images from each run was preprocessed following standard procedures including removing the first two volumes, motion correction through 6-DOF transformation to the third volume, high-pass filtering (cutoff at 100 s), slice-timing correction, spatial smoothing with 6mm full width at half maximum (FWHM), and co-registration to the structural image. Volumes with framewise displacement (FD) above 1mm were identified and censored in later analyses. Runs in which over 15% volumes were censored were excluded. An inflated brain surface of the MNI brain template was reconstructed in FreeSurfer for visualizing the result maps.

### fMRI statistical analysis

#### Action task

Functional imaging data were first analyzed using general linear models (GLM). Each run was modeled by eight experiment regressors: Three tasks (reaching, grasping, writing) and one return condition for each hand. Task phases were modeled from the onset of each trial to the return cue. Return phases were modeled from the return cue and assumed lasting one second, collapsing across actions (each hand had one return regressor). The experimental regressors were generated by convolving box-car functions spanning the periods of each condition with a standard double-gamma function. Covariates of no interests included six head motion parameters and their first derivatives, mean timeseries from the white matter and gray matter separately, and volumes with excessive head motion (FD > 1mm).

We primarily focused on the contrast of grasping versus rest that engaged wide-spread frontoparietal motor networks. To quantify hemispheric dominance, we first flipped the beta map of the grasping vs. rest contrast for each participant and then performed a Map (original, flipped) by Hand (left, right) random-effect ANOVA for each group using the FLAME1 option in FSL. The contrast of original versus flipped was then examined within this ANOVA model. All maps were thresholded at *p* < .01 and cluster-wise corrected at *p* < .05. Spherical ROIs (8mm radius) centered at the peak of each cluster were sampled, from which t-values were extracted and Map by Hand ANOVAs were performed at ROI-level (**Fig. 2B**, **Fig. 2C**). Finally, ROIs showing an interaction between Hand and Map were sampled as 8mm raduis spherical masks centered on the peak F-values (or centered at the center of gravity in the case of SMC where peak values were found in a cluster of voxels) and ANOVAs of Hand × Group were performed at ROI level (**Fig. 2C**).

To examine bimanual recruitment, we first thresholded the statistical map (of the grasping vs. rest contrast) of each hand from each participant at *p* < .01 uncorrected and obtained an overlap map between the two hands per individual. Overlap probability maps were then generated for each group by calculating percentage participants showing bimanual overlap for each voxel (**Fig. 3A**). Large-scale masks in premotor cortex, SMA, IPL, SPL, and LOC were sampled from the Juelich Atlas provided in FSL. Overlap between masks were removed from the mask with lower atlas probabilities in the overlap region. Within each mask, the proportion of voxels showing bimanual recruitment out of all voxels in the mask was calculated. The laterality index of each region was then calculated as the difference in the proportion voxel between the left and right homologou masks divided by their sum. One-way ANOVA across groups (Control, R-OBPI, L-OBPI) was then performed for each region to evaluate the effect of sensorimotor experience on the hemispheric lateralization of bimanual recruitment (**Fig. 3B**).

Finally, an Action (reaching, grasping) by Hand (left, right) random-effect ANOVA was performed for each group using the FLAME1 option in FSL. As a complement, the contrast of grasping versus reaching was also performed for each hand and each group and the overlap between hands was evaluated (**Fig. 4A**).

#### Multivariate analyses of the action task

Whole-brain searchlight multi-voxel pattern analyses (MVPA) were performed to examine representational contents associated with different actions. Preprocessed functional images with 2mm spatial smoothing were used. First, trialwise beta values were estimated in SPM12. Then a spherical ROI containing 200 voxels spanned through the volumetric brain space like a searchlight, within which a multi-voxel classification analysis was performed. Specifically, a cross-hand action decoding analysis was performed in which multi-voxel patterns of reaching and grasping from all trials of one hand were trained using a supported vector machine (SVM) and tested on trials from other hand, with the two hands switching the role across a two-fold cross-validation. The mean accuracy across the two folds was assigned to the center voxel of each searchlight, yielding a whole-brain accuracy map. Analyses were implemented in the CoSMoMVPA toolbox (Oosterhof et al., 2016).

Group-level analysis was analyzed using the Monte Carlo cluster statistics in CoSMoMVPA. At each voxel, a one-sample t-test was performed on the decoding accuracies across individuals. To evaluate significance, 100 null decoding accuracy maps were generated for each individual by shuffling the action labels across trials. These null maps were then randomly sampled to form 1000 null groups for multiple-comparisons correction at cluster level. Group-level results were thresholded at *p* < .01 and corrected at *p* < .05 (**Fig. 4B**).

#### Language task

Each run was modeled with seven experiment regressors: 4 sentence conditions (2 syntactic levels (canonical, noncanonical) by 2 semantics levels (plausible, implausible)), backward speech, responses, and prompts with no responses. The sentence and backward speech conditions were modeled as the length of each sentence/speech calculated by the number of samples divided by the sampling rate. The response condition was modeled from the beginning of the prompt to when the response was made, collapsing across sentence conditions. Probes with no response were modeled from the onset of each prompt to 4.5 sec later, assuming participants had been thinking yet failed to respond on time. Covariates of no interests were set up in the same manner as in the action task.

Language selectivity was calculated as the contrast between all sentence conditions and backward speech (**Fig. S5A**). Whole-brain comparisons of language selectivity were performed for each pair of groups (**Fig. 5D**). Results were further examined by running one-way ANOVA across all three groups at ROI level (**Fig. 5D**). A random-effect ANOVA of Syntax (canonical, noncanonical) by Semantics (plausible, implausible) was also performed and no clusters survived for either group. Therefore we primarily report results with sentence types collapsed.

Lateralization was examined by first flipping the beta map of language selectivity for each individual and contrasting the original map and the flipped map (**Fig. 5A**). Whole-brain voxel-wise comparisons were then performed between each pair of groups (**Fig. 5B**). Clusters showing significant difference were sampled and flipped to the opposite hemisphere to obtain homologous masks. For each mask, mean t-values of language selectivity (sentences vs. backward speech) and grasping activation was extracted and calculated, and then differences between the left and right homologous regions were calculated to index hemispheric dominance (**Fig. 5B**). One-way ANOVA across groups was then performed for each task (language, motor) in each region (BA44, BA45).

Finally, partial Pearson’s correlation was performed between language and motor left-lateralization (i.e. difference in t-value between the homologous ROIs in the left and right hemisphere). Language ROIs included BA44 and BA45 identified from whole-brain comparison between the Control and R-OBPI group on language lateralization (**Fig. 5B**). Motor ROIs include BA44 and frontoparietal motor areas that showed left lateralization in controls during grasping (**Fig. 2C**). Participants’ age was added as a covariate of no interest given potential effects of age on motor and language network (e.g. Olulade et al., 2020), although no correlation was found between age and the extent of left-lateralization in either tasks or ROIs.

## Data and code availability

All data and code required to reproduce all plots and statistical tests are available at https://doi.org/10.6084/m9.figshare.33421885. Any further materials will be made available upon reasonable request to the corresponding authors.

## Acknowledgements

We sincerely thank all participants with obstetric brachial plexus injury (OBPI) and their families for their participation in this study. We thank Dr. Yanchao Bi and Dr. Qing Cai for her critical comments on the manuscript. This work was supported by the National Science and Technology Innovation 2030 Major Project 2021ZD0204200 (2021ZD0204204) to Y. L., and by the National Natural Science Foundation of China (No. 82201525) to J.S.

## Author contributions

Y. L., J.S., L.C., and L.W. conecptualized the study. J.S. and L.C. recruited individuals with OBPI. Y.L. and J.S. performed experiments. Y.L., J.S., and L.Z. performed analyses. Y.L. and J.S. drafted the manuscript under the supervision of L.W.

## Declaration of interests

The authors declare no competing interests.

## Notes

### Competing Interest Statement

The authors have declared no competing interest.

