## Supplementary material for "Postnatal sensorimotor experience causally shapes hemispheric specialization across motor and language networks": SI

**Methods**

**Design of the fMRI language task**

Sentences in the language task varied in syntactic complexity and semantic plausibility. Syntactic complexity was manipulated by contrasting canonical serial-verb constructions with sentences containing embedded relative clauses and passive constructions (Skeide et al., 2014; Thibault et al., 2021. E.g. “The wolf chases the cat up the tree.” vs. “The cat was chased by the wolf and climbed up the tree.”). The latter required greater syntactic integration because listeners needed to establish hierarchical dependencies between modifiers and head nouns or resolve noncanonical thematic-role assignments. Semantic plausibility was manipulated by violating real-world event knowledge, including typical agent–patient relationships, physical constraints (e.g., body size), and social-role expectations, while keeping lexical items and syntactic structure constant (E.g. “The giraffe lowers its head and smiles at the rabbit.”

vs. “The rabbit lowers its head and smiles at the giraffe.”). Behavioral results confirmed that the manipulations were effective: across all groups, participants were slower and less accurate when processing syntactically complex or semantically implausible sentences (In an omnibus ANOVA of Group × Syntax × Semantics, main effect of Syntax: *F* (1, 284) = 14.10, *p* < .001; main effect of Semantics: F (1, 284) = 12.95, p < .001; **Fig. S11**). However, these manipulations did not elicit reliable differences in neural activation. Therefore, all sentence conditions were collapsed in the fMRI analyses presented in the main text, which focused on the general neural responses to spoken sentences.

**Behavioral analysis of the fMRI motor task**

Video-recordings of the action task were manually analyzed off-line by independent coders after reaching consensus on a subset of videos. For each trial, movement onset was determined as the frame from which the hand continued to move for five frames, and movement offset was determined as the frame from which the action is finished and no obvious adjustments were made (e.g. no re-opening and re-grasping. Subtle movements were unavoidable as the rod connecting the ball was not entirely rigid). Return offset was determined as the frame from which the hand made minimal visible movements for 5 frames. Video coders were blind to the action type of each trial.

**Articulatory representations of sentences**

To examine if the activity in BA44 during sentence listening reflected covert or implicit speech processing, we constructed representational dissimilarity matrices (RDMs) among sentences based on computationally generated articulatory trajectories (**Fig. S6B**) and correlated them with neural RDMs. Each sentence was first converted into syllable-level Hanyu Pinyin, with each syllable represented by its initial, final, and lexical tone. (**Fig. S6B**). The phonological forms were mapped to six commonly studied articulatory dimensions: labial configuration, tongue anterior-posterior position, tongue height, jaw opening, velopharyngeal configuration and tone-related F0 (Wu & Shih, 2009; Lee & Zee, 2003; Tiede et al., 2019; Shaw et al., 2016; Chang et al., 2016; Chiu & Lu, 2021; Moisik et al., 2014; Huang & Liao, 2017). These phonological parameters were converted into time-varying target trajectories by blending initial and final weights via a raised-cosine function within each syllable, incorporating the interpolated tonal contour, and concatenating the resulting frame-wise vectors across the sentence (Saltzman 2003; Parrell et al., 2013). The direction and relative ordering of these dimensions were defined based on established phonetic descriptions and articulatory evidence for Mandarin (**Appendix B**). Varying the exact numeric values while preserving the theoretically defined ordinal relationships yielded highly similar articulatory RDMs (mean Pearson’s r = 0.902, SD = 0.018) and therefore our main conclusions remain robust to exact parameter choices.

To model the temporal evolution of these articulatory states during speech production, the normalized targets were converted into continuous articulatory trajectories using a critically damped second-order task-dynamic system (**Eq. (1)**, **Fig. S6B;** Saltzman & Munhall, 1989; Browman & Goldstein, 1992; Nam et al., 2012). This formulation models each articulatory dimension as a dynamic state that smoothly approaches its time-varying target while maintaining temporal continuity across syllable boundaries.

|  | $\ddot{x}+2\zeta\omega\dot{x} + \omega^{2}(\dot{x} -g)$ = 0 | (1) |
| --- | --- | --- |

Where *g* is the articulatory target, *x* is the dynamic trajectory, and $\text{ζ}$ is the damping ratio, which controls how rapidly the articulatory trajectory settles toward the articulatory target. It was set to 1 to achieve a fast and non-oscillatory response (Saltzman & Munhall, 1989; Kuberski & Gafos, 2019).

To quantify similarity between sentences, the resulting six-dimensional trajectories were aligned using multivariate dynamic time warping (DTW) that accounts for differences in sentence length and temporal alignment while preserving the multidimensional structure of the trajectories (Sakoe & Chiba, 1978; Shokoohi-Yekta et al., 2017). All six dimensions were jointly aligned using a common warping path, and the cumulative DTW cost was normalized by path length to obtain the mean local distance along the optimal alignment path.

We note that without directly collecting articulatory recordings such as electromagnetic articulography (EMA) data, this model provides a theory-driven approximation of articulatory dynamics.To assess whether the articulatory model captured the expected structure of the speech materials. Our design involved manipulating the semantic plausibility (plausible, implausible) and syntactic complexity (canonical, noncanonical) of each context sentence (**Method**). As a result, each context contained four sentences that share the same lexical items but differed in syntactic structures. For example, “A monkey is riding on an elephant and picking apples” and “An elephant is riding on a monkey and picking apples”. Because sentences within the same context shared the same lexical content, we expected greater articulatory similarity for sentence pairs within the same context than for those across contexts. To test this prediction, we performed a permutation test by randomly shuffling the context labels across sentences and calculating the difference between overall within-context and across-context similarity to generate a null distribution. Consistent with this prediction, articulatory similarity was significantly higher for within- than across-context sentence pairs (permutation *p* < .001; **Fig. S6B**), suggesting that our articulatory model to capture the expected structure of the speech materials.

Finally, neural RDMs were constructed by first estimating the beta value during sentence listening in each trial using SPM12. Beta values within the BA44 ROI were then sampled and Euclidean distances between sentences were calculated, forming an RDM across sentences for each participant. The neural RDM from each participant was correlated with the articulatory RDM using Pearson’s correlation.

**Supplementary Figures**

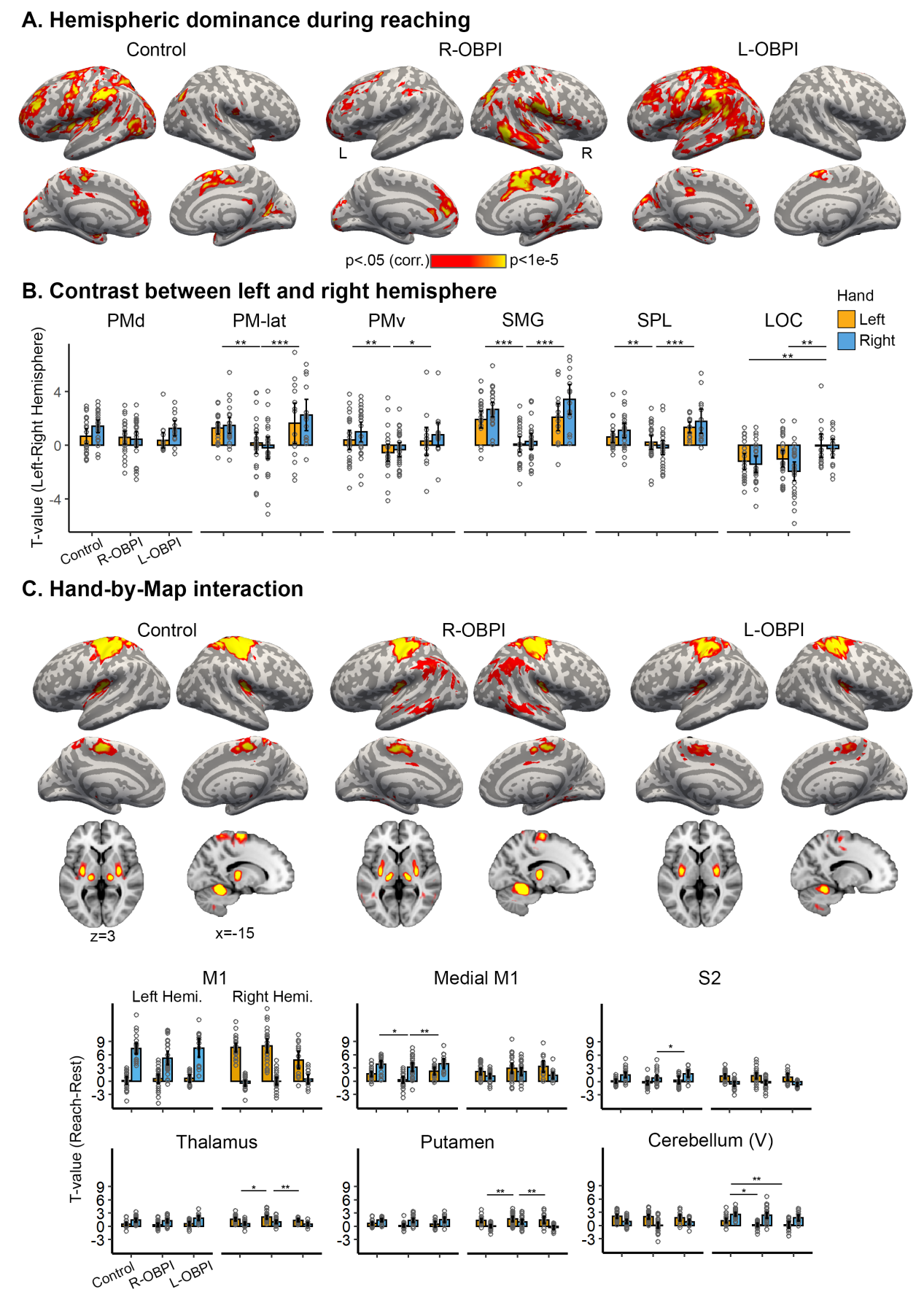

**Fig. S1. Reaching actions show similar hemispheric dominance patterns as with grasping.**

(A) Hemispheric dominance maps during reaching. Consistent with grasping, controls showed pronounced left-hemisphere dominance across premotor, parietal, and lateral occipital cortices, while sparing the central sulcus (CS). This leftward dominance was substantially reduced in individuals with right-sided obstetric brachial plexus injury (R-OBPI), whereas individuals with left-sided OBPI (L-OBPI) exhibited dominance patterns comparable to controls. Statistical maps show the contrast between original vs. flipped maps collapsed across hands (cluster-wise corrected at *p* < 0.05).

(B) Hemispheric dominance analysis at ROI level. The same ROIs were used as in the analyses of grasping (**Fig. 2B**). Similar to results from grasping, R-OBPI participants exhibited significantly reduced left-hemisphere dominance compared with both controls and L-OBPI participants in lateral PMd, PMv, SMG and LOC, independent of the hand used for action. Error bars denote 95% CI. Circles indicate individual participants. Asterisks denote significant post hoc comparisons. * p < .05; ** p < .01; *** p < .001. Raw activation levels for each hand in each hemisphere are shown in **Fig. S2**.

(C) Hemisphere-by-hand interaction reveals hand-dependent hemispheric dominance that is stable across groups. These regions, including primary sensorimotor cortex (SMC), secondary somatosensory cortex (S2), medial primary motor cortex (M1-medial), thalamus, and putamen, exhibited the expected contralateral organization. Cerebellar lobule V showed ipsilateral organization.

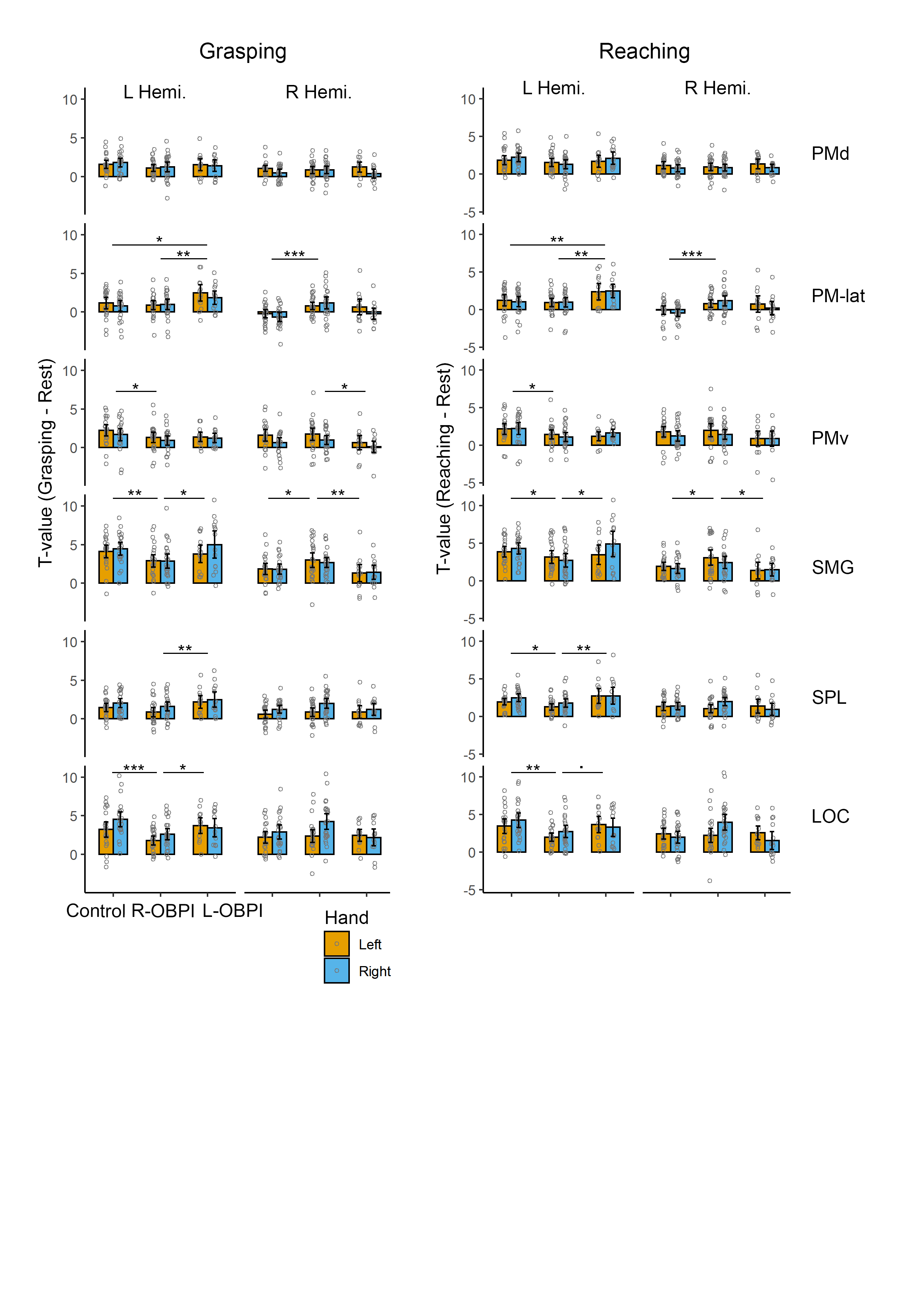

**Fig. S2.** Decomposing activation of each hand in bilateral ROIs that showed a main effect of Map (**Fig. 2A and Fig. S1A**), for grasping (left) and reaching (right). There is a general trend that individuals with R-OBPI showed decreased activation in the left hemisphere and increased activation in the right hemisphere, resulting in the general reduction in left-hemisphere dominance.

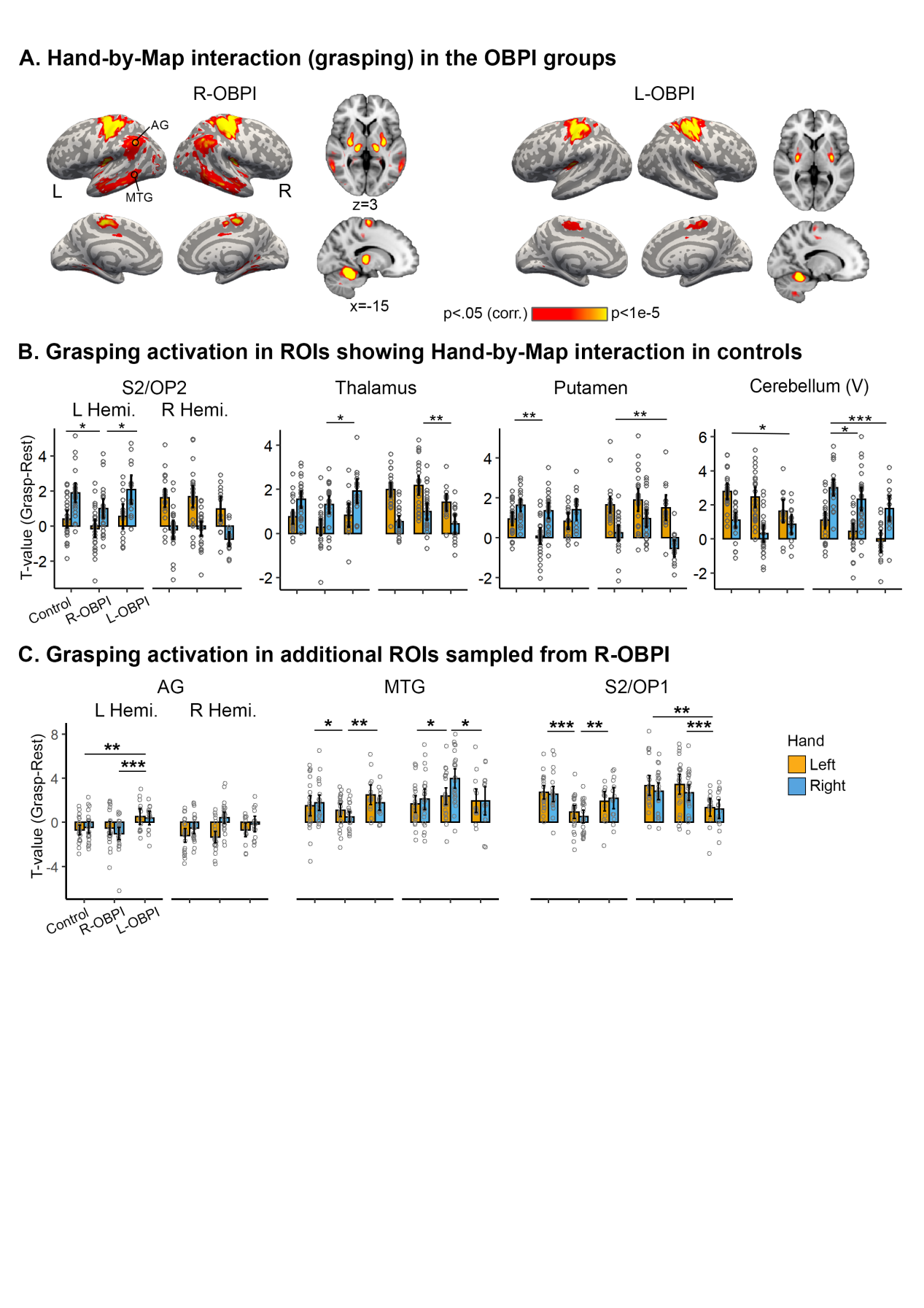

**Fig. S3. Hand-by-Map interaction during grasping actions (complementing Fig. 2C)**

(A) Whole-brain map showing Hand-by-Map interaction for the grasping action in the OBPI groups, complementing **Fig. 2**. Similar patterns were found in core regions as with controls. R-OBPI additionally showed interaction in angular gyrus (AG) and middle temporal gyrus (MTG).

(B) More ROIs that show a Hand × Map interaction (complementing **Fig. 2C**). Motor activation of each hand in each hemisphere of thalamus and putamen showed contralateral representation across groups; Cerebellum (lobule V) exhibited the canonical ipsilateral representation.

(C) Grasping activation of each hand in AG, MTG, and S2/Central Operculum (OP2) in R-OBPI.

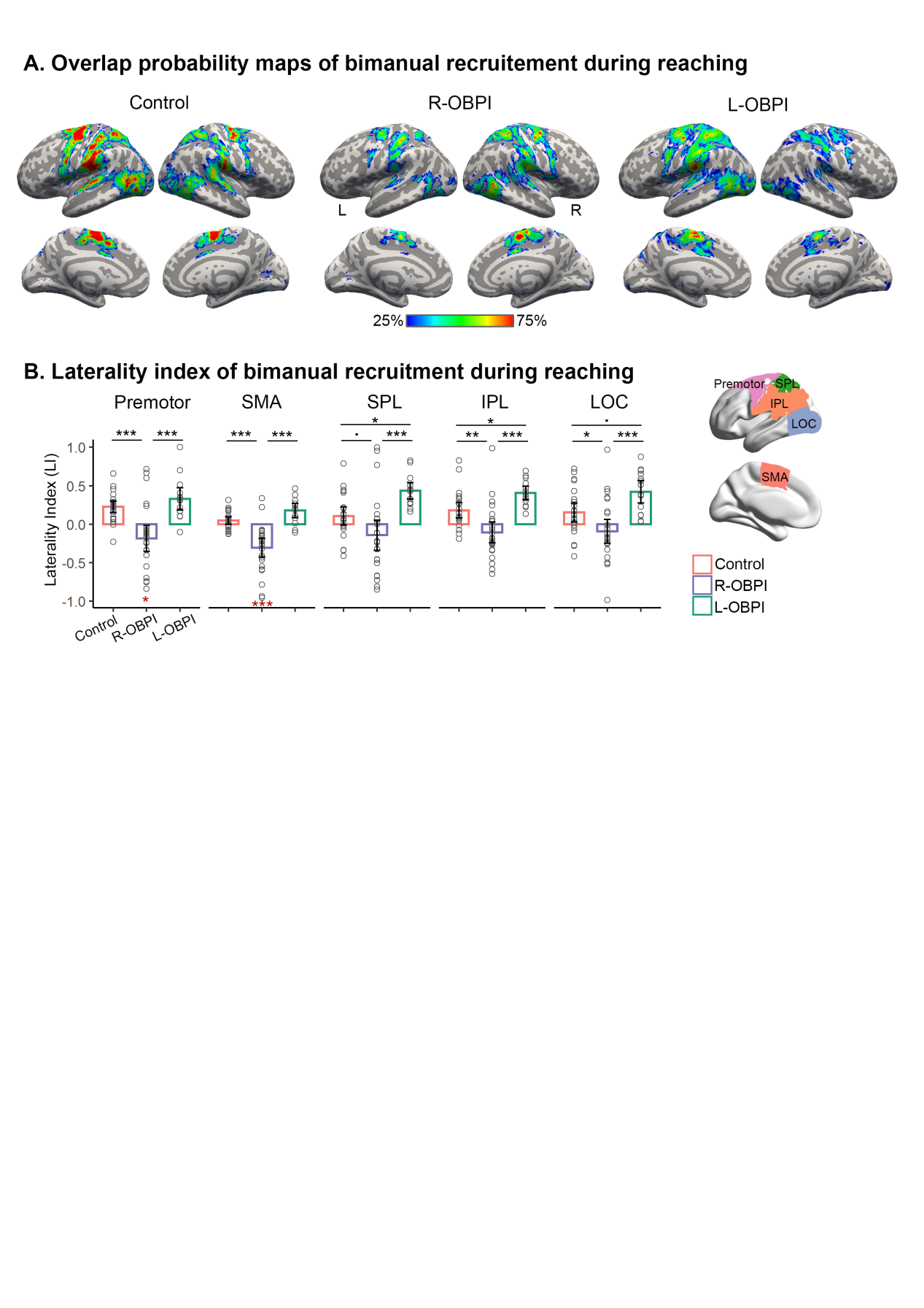

**Fig. S4.** **Experience-dependent reorganization of bimanual recruitment in higher-order motor regions during reaching actions**

**(A)** Overlap probability maps of bimanual recruitment for each group for the reaching action. Consistent with grasping, in Controls and the L-OBPI group, bimanual recruitment was concentrated in left premotor, parietal, and lateral occipital cortices. In contrast, the R-OBPI group exhibited a larger spatial extent and greater inter-individual overlap of bimanual recruitment in the right hemisphere.

**(B)** Laterality indices (LI) of bimanual recruitment in bilateral premotor cortex, supplementary motor area (SMA), superior parietal lobule (SPL), inferior parietal lobule (IPL), and lateral occipital cortex (LOC). Positive values indicate left-hemisphere dominance. Relative to Controls and the L-OBPI group, the R-OBPI group showed significantly reduced or reversed lateralization, with significant right-hemisphere dominance in premotor cortex and SMA (red asterisks). Conversely, the L-OBPI group exhibited enhanced left lateralization in SPL, IPL, and LOC compared with Controls. Error bars denote 95% CI. * p < .05; ** p < .01; *** p < .001.

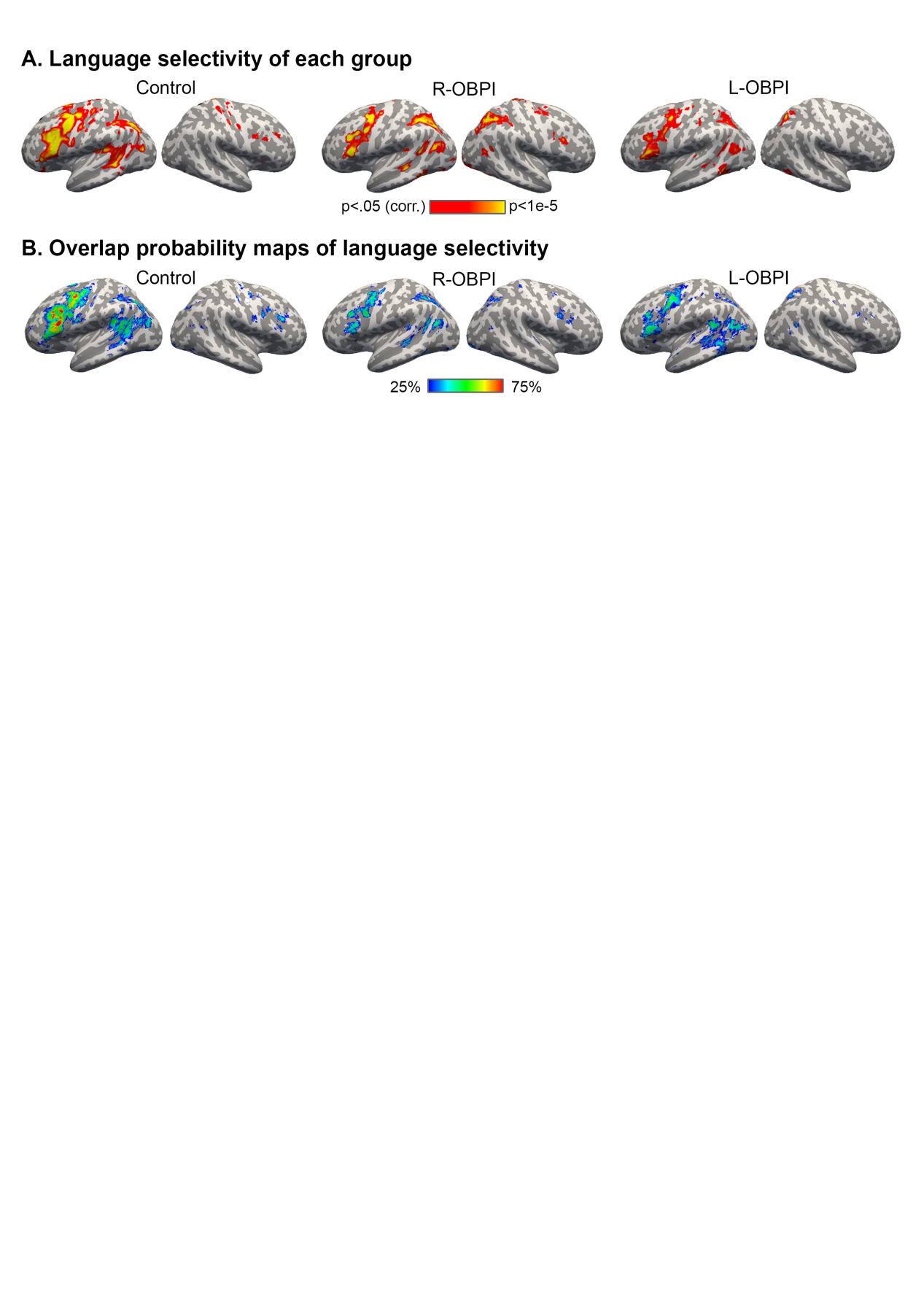

**Fig. S5. (A)** Language selectivity maps of language processing obtained from the voxel-wise mirror subtraction analysis. All groups exhibited language-selective responses within a predominantly left-lateralized frontotemporal network encompassing inferior and middle frontal gyri and middle/posterior temporal cortices. **(B)** Overlap probability maps of language selectivity in each group.

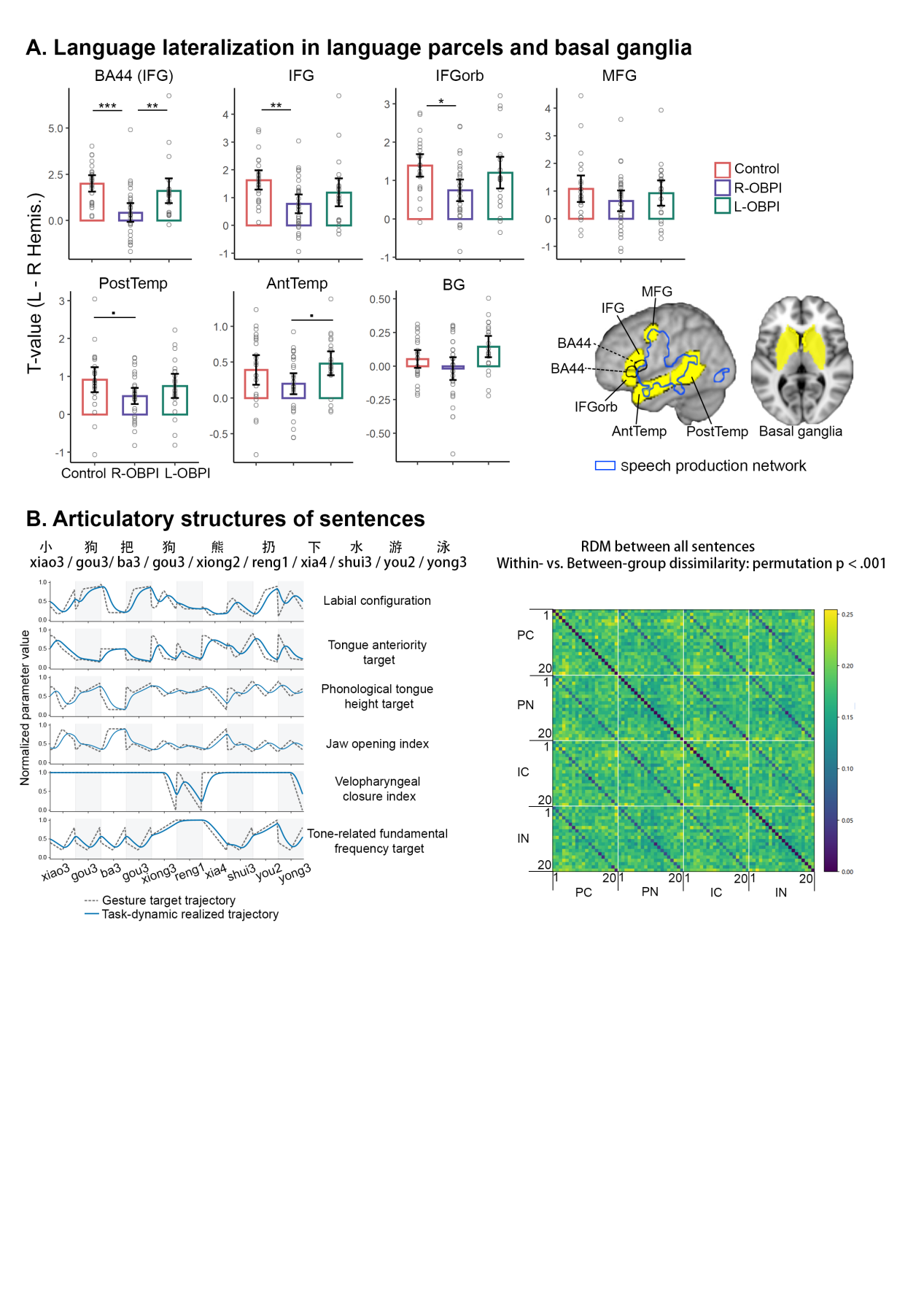

**Fig. S6. (A)** Language hemispheric dominance in language parcels from Fedorenko et al., (2010), including IFG, IFGorb, MFG, anterior temporal lobe, and posterior temporal lobe. The intersection between BA44 from **Fig. 5B** and IFG (BA44 (IFG)) and ROIs of bilateral basal ganglia sampled based on Harvard-Oxford Subcortical Structural Atlas was also included.

**(B)** Dynamic articulatory trajectories of each articulatory domain for an example sentence (shown in Chinese characters and in Hanyu Pinyin). The RDM on the right shows the dissimilarity structure across all 80 sentences used in the experiment. Numbers along the axes denote context numbers, with sentences sharing context numbers share lexical items; blocks denote semantic and syntactic variations of the sentences (PC: Semantically plausible and Syntactically canonical; PN: Plausible-Noncanonical; IC: Implausible-Canonical; IN: Implausible-Noncanonical).The off-diagonal bands parallel to the main diagonal denote lower dissimilarity between sentences within the same context vs. across contexts (permutation *p* < .001).

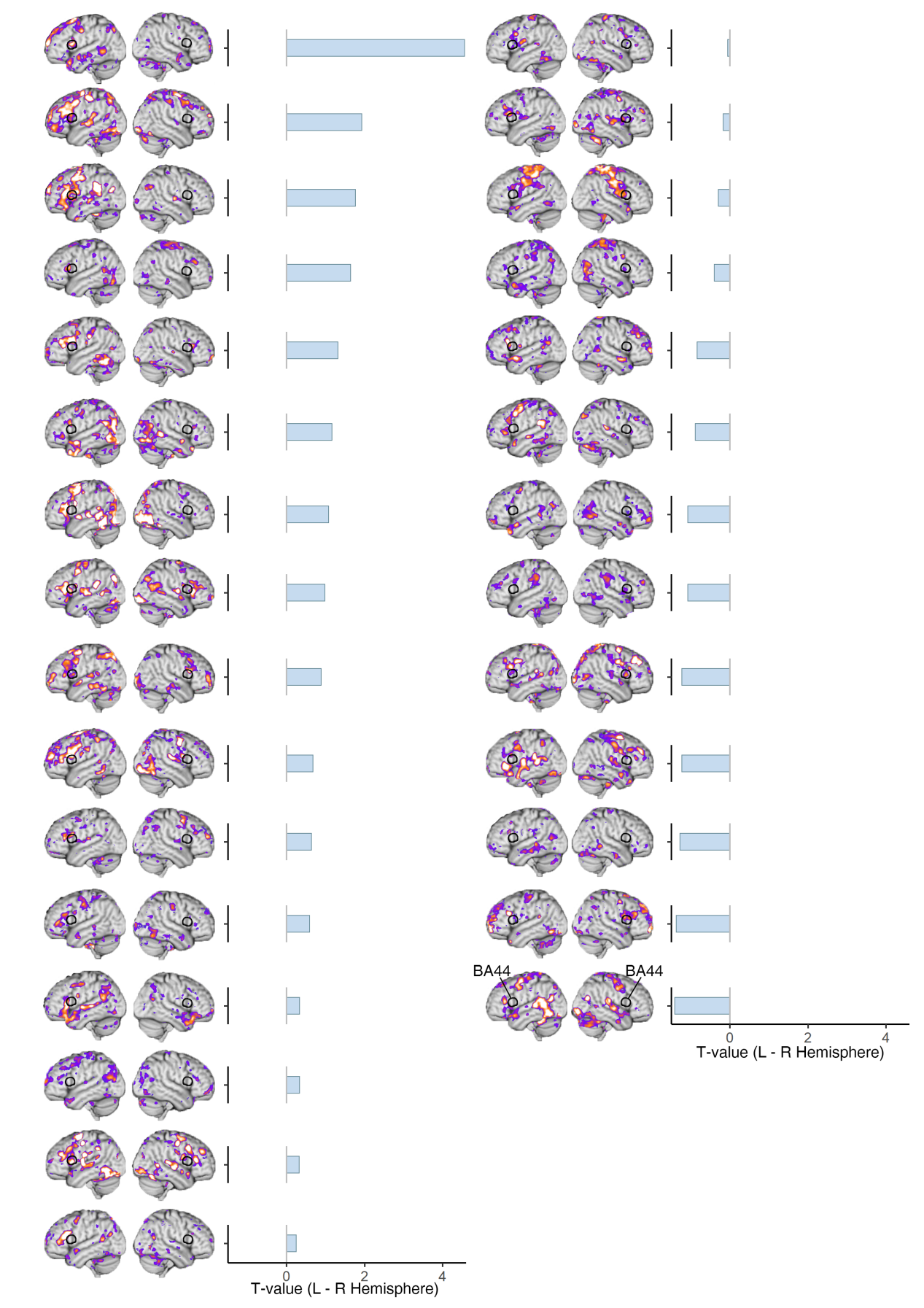

**Fig. S7.** Individual language selectivity maps of the R-OBPI group. Maps are ordered based on the hemispheric dominance (left - right hemisphere) in left BA44 sampled from the contrast of Control vs. R-OBPI in terms of language hemispheric dominance (bar graphs; **Fig. 5B**). 44.8% of participants showed right-hemisphere dominance (the right column).

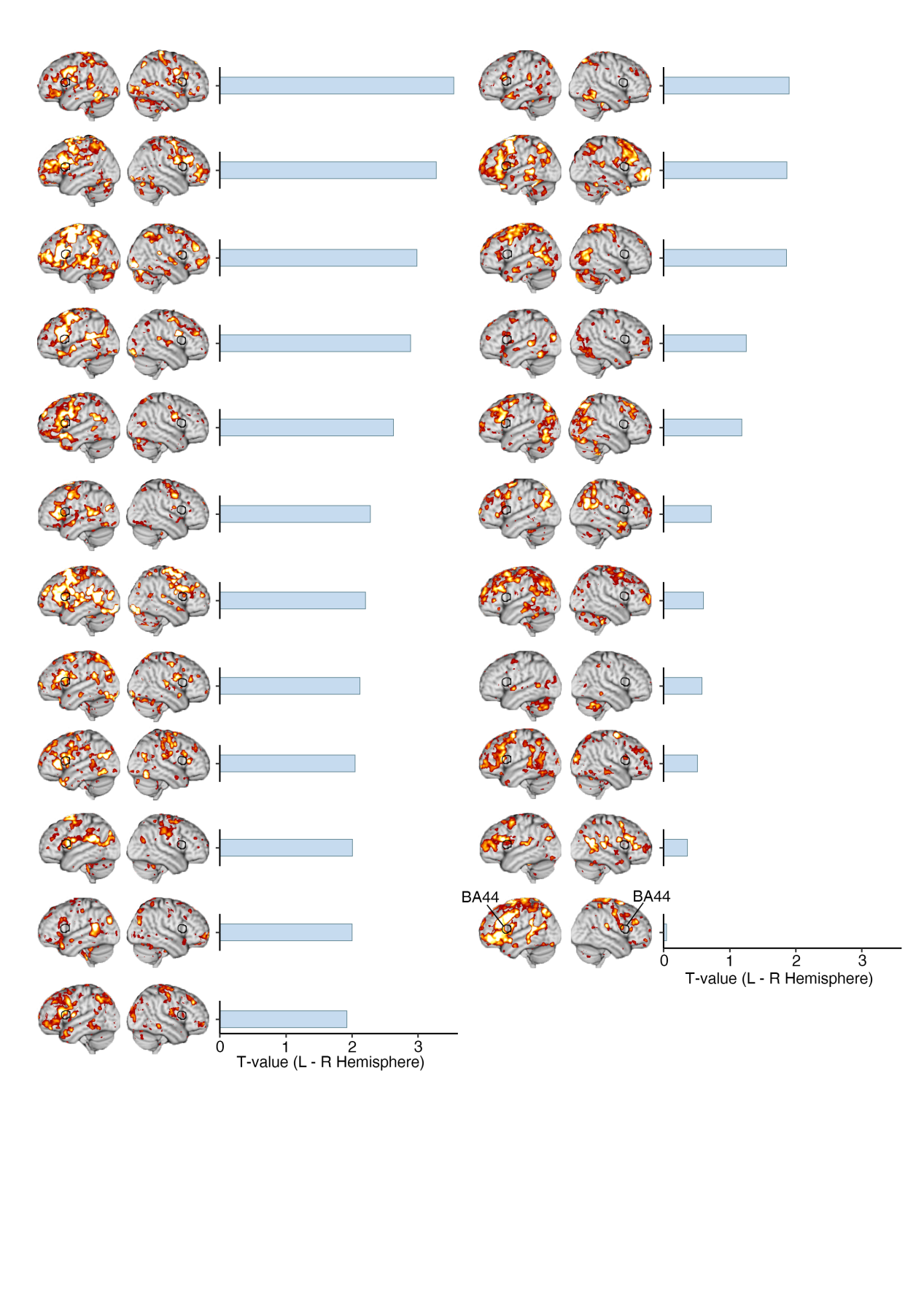

**Fig. S8.** Individual language selectivity maps of the Control group. Maps are ordered based on the hemispheric dominance (left - right hemisphere) in left BA44 sampled from the contrast of Control vs. R-OBPI in terms of language hemispheric dominance (bar graphs; **Fig. 5B**). All participants showed left-hemisphere dominance.

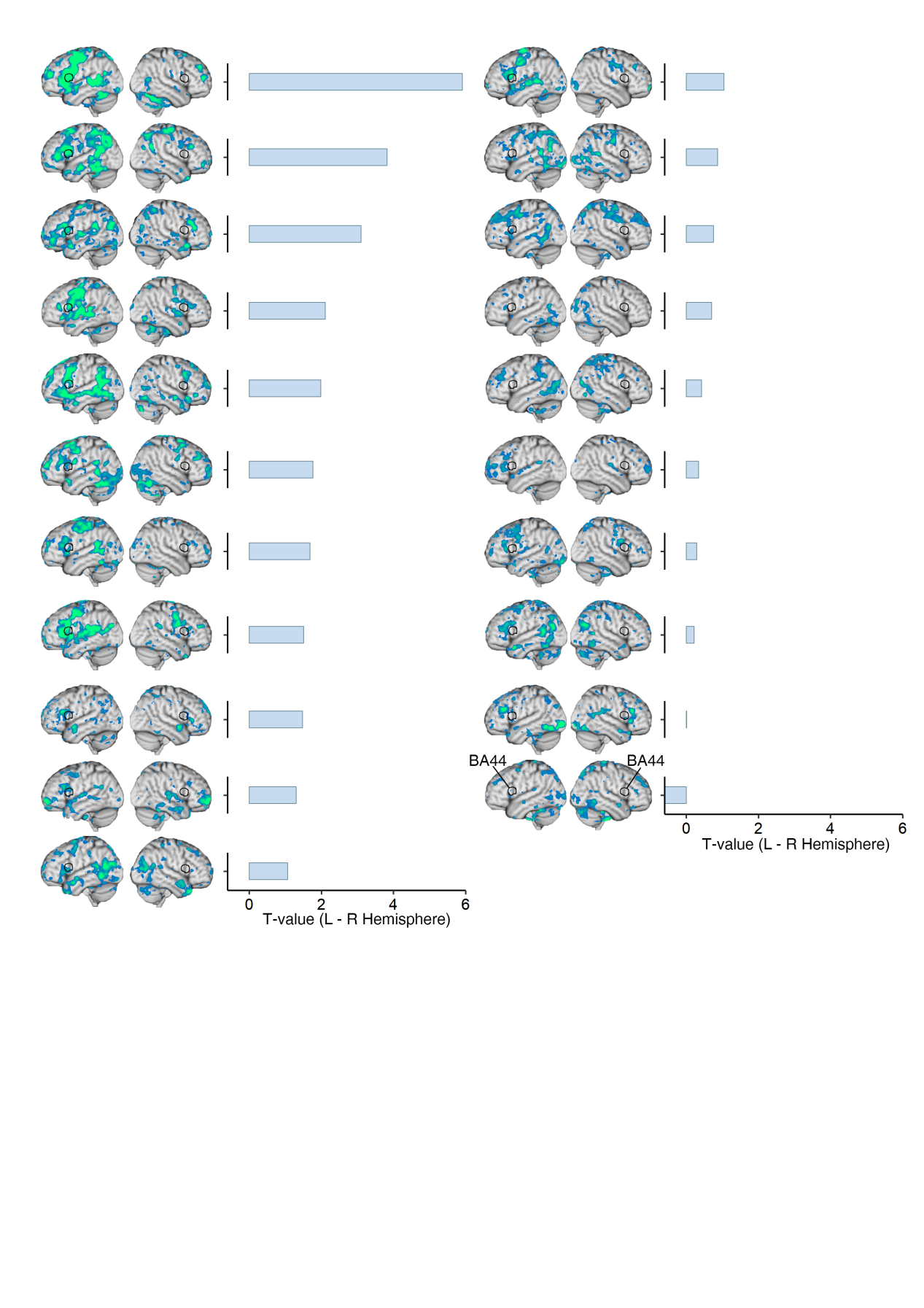
**Fig. S9.** Individual language selectivity maps of the L-OBPI group. Maps are ordered based on the hemispheric dominance (left - right hemisphere) in left BA44 sampled from the contrast of Control vs. R-OBPI in terms of language hemispheric dominance (bar graphs; **Fig. 5B**). All but one participants showed left-hemisphere dominance.

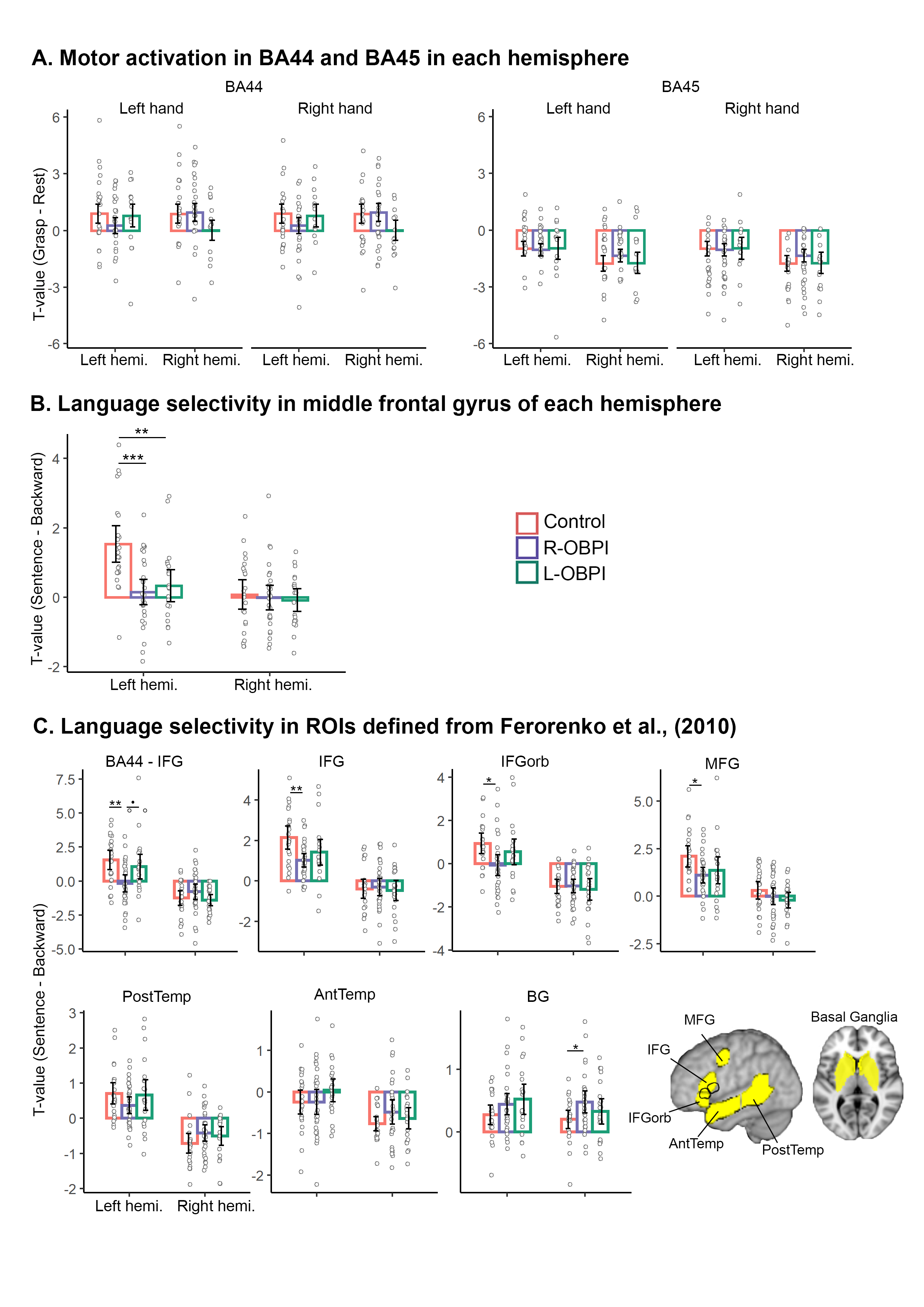

**Fig. S10.** Motor and language response profiles in language ROIs.

(A) Activation in BA44 and BA45 (as in **Fig. 5B**) during grasping actions. Whereas BA44 responded to the motor task, BA45 was not activated.

(B) Language selectivity in middle frontal gyrus (MFG) in each hemisphere, with left MFG sampled from the whole-brain contrast betwen Control and R-OBPI in language selectivity (**Fig. 5D**). Right MFG was obtained by mirror-flipping the left MFG. Both OBPI groups showed reduced language selectivity in BA44 compared to controls.

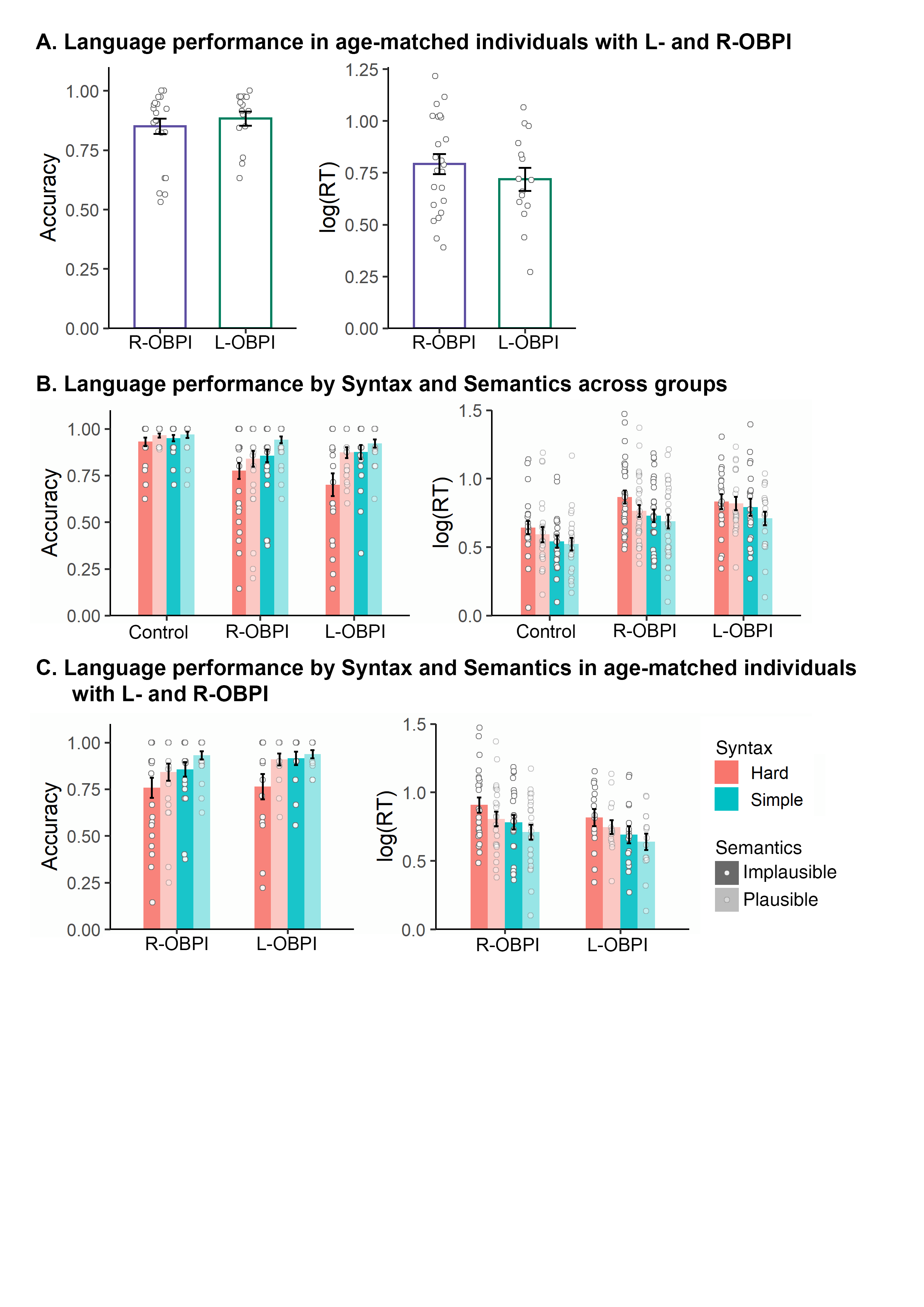

**Fig. S11. Behavioral performance during the fMRI language task.**

(A) No difference in accuracy (*t* (36) = 0.721, *p* = .476) or reaction time (*t* (36) = 0.994, *p* = .327) was found between age-matched individuals with L- and R-OBPI.

(B) Accuracy and reaction time as a function of syntactic difficulty and semantic plausibility. For accuracy, there was a main effect of Group, *F* (2, 284) = 12.74, *p* < .001, a main effect of Syntax, *F* (1, 284) = 14.10, *p* < .001, and main effect of Semantics, *F* (1, 284) = 12.95, *p* < .001. Both OBPI groups made more mistakes than controls (post-hoc *p*s < .001), with no difference between the OBPI groups (post-hoc *p* = .908). No interaction effects were found. Similarly, for log-transformed reaction time, there was a main effect of Group, *F* (2,284) = 20.46, *p* < .001), with both OBPI groups responded more slowly than controls (post-hoc *p*s < .001) and no differences between the OBPI groups (post-hoc *p* = .734). The main effect of Syntax was significant, *F* (1, 284) = 10.03, *p =* .002), and the main effect of Semantics was marginal, *F* (1, 284) = 3.39, *p* = .067). No interaction effects were found.

(C) No group difference was found between age-matched individuals with L- and R-OBPI (main effect of Group: *F* (1, 144) = 1.22, *p* = .272). No interaction effects were found between Group and either syntactic difficulty or semantic plausibility (*p*s > .3).

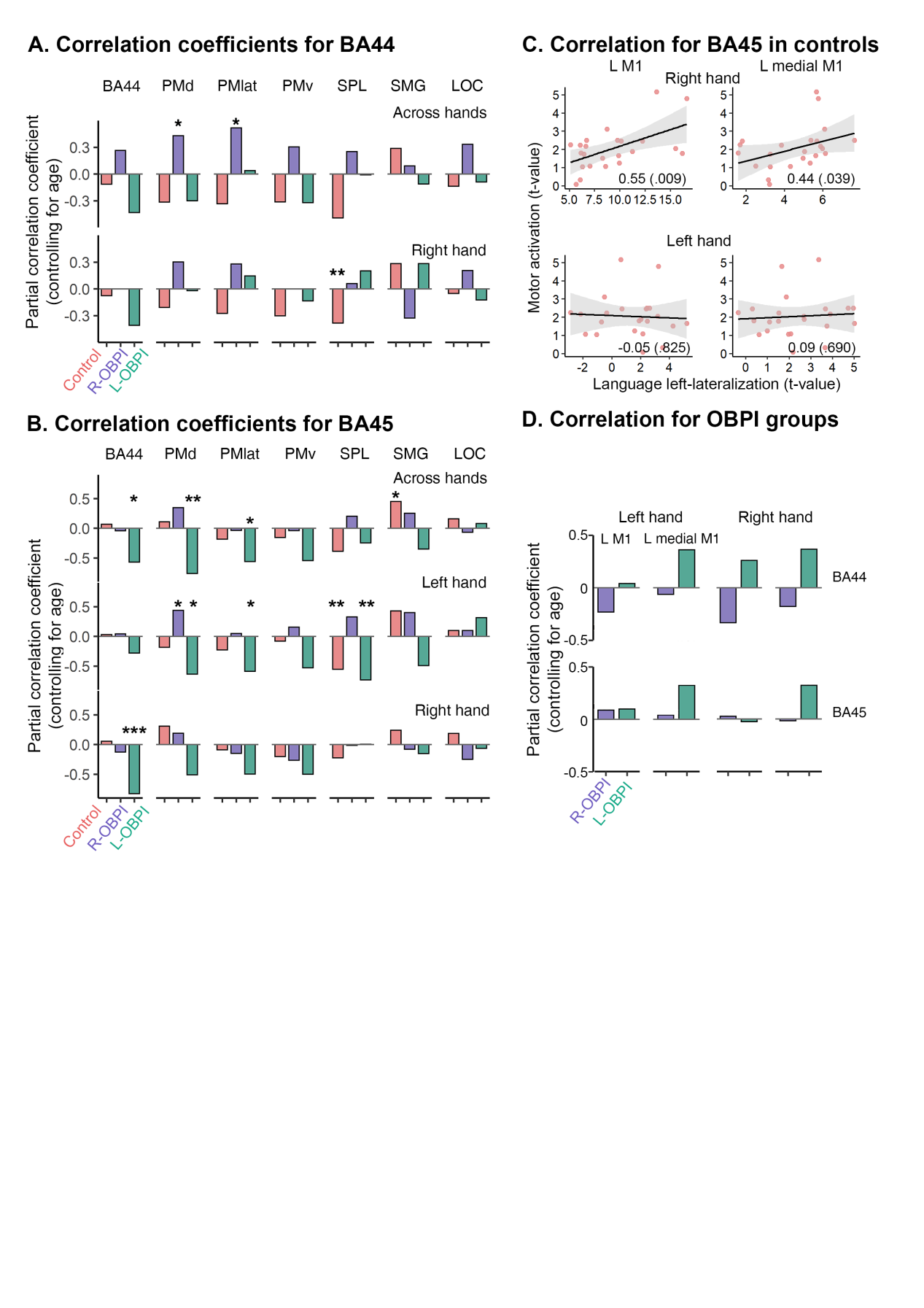

**Fig. S12.** Correlation between language and motor networks.

(A) Correlation between language left-lateralization in BA44 and motor left-lateralization in BA44 and frontoparietal regions, with motor left-lateralization calculated as the average across hands or for the right hand. Individuals with R-OBPI showed positive correlation in left PMd and lateral premotor cortex, which were primarily driven by the left hand (**Fig. 6**). Controls and individuals with L-OBPI primarily showed weak or negative correlation.

(B) Correlation between language left-lateralization in BA45 and motor left-lateralization in BA44 and frontoparietal regions, with motor left-lateralization calcuated as the average across hands and for each hand separately.

(C) Correlation between language left-lateralization in BA45 and motor activation (during grasping) in left SMC and left medial M1. As with BA44, correlation was found specifically for right-hand movements.

(D) Correlation coefficients between language left-lateralization in BA44 (upper) and BA45 (lower) with motor activation (during grasping) in left SMC and left medial M1, in the OBPI group. Unlike patterns in controls, no correlation was found in the OBPI groups.

**Appendix A. Clinical characteristics and surgical history of participants with obstetric brachial plexus injury (OBPI)**

| ID | Sex | Age at test | Affected side | Age at operation (mo.) | Preope. Narakas classification | Intraope. injury type | Surgery |
| --- | --- | --- | --- | --- | --- | --- | --- |
| E01 | M | 11 | L | 4 | 4 | C5R, C6-T1A | C5→ MC (with grafting), SAN→SSN, ICN3-5→MCN |
| E02 | M | 22 | R | 4 | 4 | —— | Brachial plexus neurolysis |
| E03 | M | 20 | R | 3 | 4 | C5-C6R, C7-T1A | C5→ ADUT&PDUT (with grafting), C6→ ADUT&MC (with grafting), SAN→SSN |
| E04 | M | 11 | R | 4 | 4 | C5-C6R, C7-C8A | C5→UT&C8 (with grafting), C6→C7&C8 (with grafting), SAN→SSN |
| E05 | M | 12 | L | 3 | 4 | —— | —— |
| E06 | M | 12 | R | 3 | 4 | C5R, C6-T1A | C5→ MC (with grafting), SAN→SSN, ICN3-5→MCN, ICN3 sensory branch→lateral root of MN |
| E07 | M | 8.5 | L | 4 | 4 | C5-C6R, C7-T1A | C5→ MC (with grafting), C6→C7&PDLT (with grafting), SAN→SSN, ICN3-5→MCN |
| E08 | M | 16 | R | 7 | 2 | C5A, C6R, C7A | C6&partial C5→PC (with grafting), ICN3-6→MCN, SAN→SSN |
| E09 | F | 15 | R | 4 | 4 | C5-C6R, C7-C8A, T1 partial A | C5→ ADUT&PDUT (with grafting), C6→MT&C8 (with grafting), SAN→SSN |
| E10 | F | 7.6 | L | 4 | 4 | C5R, C6 partial A, C7-T1A | C5→ MC (with grafting), C6→C7&ADUT (with grafting), SAN→SSN, ICN8-10→MCN |
| E11 | M | 17 | R | 6 | 4 | —— | —— |
| E12 | M | 14 | L | 7 | 4 | C5-C6R, C7-C8A | C5→UT&PDUT (with grafting), C6→C7&C8 (with grafting), SAN→SSN |
| E13 | M | 14 | L | 7 | 2 | C5-C6R, C7A | C5→PC (with grafting), C6→LC (with grafting), SAN→SSN |
| E14 | M | 13 | R | 4 | 2 | C5-C7A | SAN→SSN, ICN3-5→MCN |
| E15 | F | 8.4 | L | 4 | 4 | —— | —— |
| E16 | M | 8.3 | L | 3 | 2 | C5-C6R, C7A | C5→ ADUT&SSN (with grafting), C6→MT&PDUT (with grafting) |
| E17 | M | 8.3 | R | 3 | 2 | —— | —— |
| E18 | F | 15 | R | 4 | 4 | C5-C7R, C8-T1A | C5→ MC (with grafting), C6→C7 (with grafting), SAN→SSN, ICN3-6→MCN |
| E19 | F | 11 | R | 3 | 4 | C5-C7R, C8-T1A | C5→MT&PDLT (with grafting), C6→ MC (with grafting), C7→ ADUT (with grafting), SAN→SSN, ICN7&9-10→MCN |
| E20 | M | 12 | L | 6 | 2 | C5-C6R, C7A | C5→ ADUT&SSN (with grafting), C6→MT& PDUT (with grafting) |
| E21 | M | 11 | R | 7 | 2 | C5-C6R, C7A | C5→UT (with grafting), C6→MT (with grafting), SAN→SSN |
| E22 | M | 10 | L | 3 | 4 | C5-C6R, C7-C8A | C5→C8 (with grafting), C6→PDMT (with grafting), SAN→SSN, ICN4-6→MCN |
| E23 | F | 11 | L | 3 | 4 | C5-C6R, C7-T1A | C5→PDLT&C7 (with grafting), C6→ MC (with grafting), SAN→SSN, ICN7-9→MCN |
| E24 | M | 11 | L | 5 | 4 | C5R, C6-T1A | C5→ MC (with grafting), SAN→SSN, ICN3-5→MCN |
| E25 | M | 13 | L | 7 | 4 | C5-C6R, C7-C8A, T1 partial A | C5→ ADUT& PDUT (with grafting), SAN→SSN, C6→C7&C8 (with grafting) |
| E26 | F | 20 | R | 4 | 2 | C5-C6R, C7A | C5→UT (with grafting), C6→MT (with grafting), SAN→SSN |
| E27 | M | 15 | R | 6 | 2 | C5-C7A | SAN→SSN, ICN3-6→MCN |
| E28 | M | 21 | R | 6 | 2 | C5-C6R, C7A | C5→ PDUT&MT (with grafting), C6→ ADUT&MT (with grafting), SAN→SSN |
| E29 | M | 12 | R | 3 | 2 | C5-7A | SAN→SSN, ICN3-5→MCN |
| E31 | F | 13 | R | 5 | 4 | C5-C6R, C7-C8A, T1 partial A | C5→ ADUT&C8 (with grafting), C6→C7&C8 (with grafting), SAN→SSN |
| E33 | F | 9.4 | L | 3 | 2 | C5-C6R, C7A | C5→ ADUT&SSN (with grafting), C6→MT& PDUT (with grafting) |
| E34 | M | 8.5 | L | 3 | 4 | C5R, C6-T1A | C5→ MC (with grafting), SAN→SSN, ICN3-5→MCN |
| E35 | M | 19 | L | 4-5 year | 2 | —— | Brachial plexus neurolysis |
| E36 | M | 7.9 | L | 3 | 2 | C5R, C6-C7A | C5→MT (with grafting), SAN→SSN, ICN8-10→MCN |
| E37 | M | 16 | R | 3 | 4 | C5-C6R, C7-C8A | C5→ ADUT&PDUT (with grafting), SAN→SSN, C6→C7&C8 (with grafting) |
| E38 | M | 16 | R | 3 | 4 | C5-C6R, C7-C8A, T1 partial A | C5→PC, C6→ MC, SAN→SSN, ICN3-5→MCN |
| E39 | M | 18 | R | 3 | 2 | C5-C7R | C5→UT&C8 (with grafting), C6→ ADUT&PDUT (with grafting), C7→MT& PDUT (with grafting), SAN→SSN |
| E40 | F | 16 | L | 3 | 4 | C5-C6R, C7-T1A | C5→ MC (with grafting), C6→C7&PDLT (with grafting), SAN→SSN |
| E41 | M | 15 | L | 5 | 4 | C5-C6R, C7-C8A, T1 partial A | C5→C7&PDUT (with grafting), C6→C7&C8 (with grafting), SAN→SSN, ICN3-5→MCN |
| E42 | M | 10 | L | 4 | 2 | C5R, C6-C7A | C5→C7&PDLT (with grafting), SAN→SSN, ICN3-5→MCN |
| E43 | F | 17 | R | 5 | 4 | C5R, C6-C8A, T1 partial A | C5→C7&C8 (with grafting), SAN→SSN, ICN3-6→LC |
| E44 | F | 11 | R | 3 | 4 | C5R, C6-T1A | C5→ MC (with grafting), SAN→SSN, ICN6-8→MCN |
| E45 | F | 15 | R | 3 | 4 | C5-C6R，C7-T1A | C5→C7&PDLT (with grafting), C6→ MC (with grafting), SAN→SSN, ICN3-5→MCN |
| E46 | M | 9.6 | R | 3 | 4 | C5R, C6-T1A | C5→ MC (with grafting), SAN→SSN, ICN3-5→MCN, ICN3 sensory branch→lateral root of MN |
| E47 | F | 12 | R | 3 | 2 | C5-C6R, C7A | C5→ ADUT&SSN (with grafting), C6→MT& PDUT (with grafting) |
| E48 | M | 13 | L | 3 | 2 | C5-C6R, C7A | C5→RN (with grafting), SAN→SSN (with grafting), ICN3-5→MCN |
| E49 | F | 17 | L | 5 | 2 | C5-C6R, C7A | C5→ PDUT&MT (with grafting), C6→MT& PDUT&ADUT (with grafting) |
| E50 | F | 12 | R | 7 | 2 | C5-C6R, C7A | C5→ ADUT&SSN (with grafting), C6→MT& PDUT (with grafting) |
| E51 | M | 10 | R | 5 | 4 | C5-C6R, C7-C8A, T1 partial A | C5→C8 (with grafting), C6→MT& PDUT (with grafting), SAN→SSN, ICN3-5→MCN |
| E52 | F | 11 | R | 3 | 2 | C5-C6R, C7A | C5→ PDUT (with grafting), SAN→SSN (with grafting), C6→ ADUT (with grafting) |
| E53 | M | 12 | R | 3 | 4 | C5R, C6-T1A | C5→ MC (with grafting), SAN→SSN, ICN6-8→MCN |
| E54 | M | 21 | R | 3 | 4 | C5-T1A | SAN→SSN, contralateral C7→MCN |

Intraope. injury type: R - Rupture, A - Avulsion

Surgery: ADUT, Anterior Division of Upper Trunk; ICN, intercostal Nerve; LC, Lateral Cord; MC, Medial Cord; MCN, Musculocutaneous Nerve; MN, Median nerve; MT, Middle Trunk; PC, Posterior Cord; PDMT, Posterior Division of Middle Trunk; PDLT, Posterior Division of Lower Trunk; PDUT, Posterior Divison of Upper Trunk; RN, Radial Nerve; SAN, Spinal Accesory Nerve; SSN, Suprascapular Nerve; UT, Upper Trunk;

**Appendix B. Mapping from Hanyu Pinyin to articulatory target parameters**

| Sound | L | Tx | Ty | J | V |
| --- | --- | --- | --- | --- | --- |
| b, p | .95 | .50 | .50 | .35 | 1 |
| m | .95 | .50 | .50 | .35 | 0 |
| f | .65 | .45 | .50 | .35 | 1 |
| d, t | .45 | .80 | .50 | .35 | 1 |
| n | .45 | .80 | .50 | .35 | 0 |
| l | .45 | .80 | .60 | .35 | 1 |
| z, c, s | .40 | .85 | .55 | .35 | 1 |
| zh, ch, sh, r | .45 | .65 | .70 | .35 | 1 |
| j, q, x | .35 | .85 | .75 | .35 | 1 |
| g, k, h | .30 | .20 | .75 | .35 | 1 |
| a | .20 | .50 | .15 | .90 | 1 |
| o | .80 | .25 | .50 | .60 | 1 |
| e | .30 | .35 | .50 | .55 | 1 |
| i | .10 | .90 | .90 | .30 | 1 |
| u | .90 | .15 | .85 | .30 | 1 |
| ü | .90 | .85 | .90 | .30 | 1 |
| I (zi, ci, si) | .20 | .85 | .58 | .35 | 1 |
| I (zhi, chi, shi, ri) | .30 | .65 | .70 | .35 | 1 |
| n | .45 | .80 | .50 | .35 | 0 |
| ng | .30 | .20 | .70 | .35 | 0 |

L:Labial configuration; Tx:Tongue anterior-posterior position; Ty:Tongue height; J: Taw opening; V: Velopharyngeal configuration.
